# A phosphoregulated RhoGEF feedback loop tunes cortical flow driven amoeboid migration *in vivo*

**DOI:** 10.1101/2021.10.29.466358

**Authors:** Benjamin Lin, Jonathan Luo, Ruth Lehmann

## Abstract

Cortical flow driven amoeboid migration utilizes friction from retrograde cortical actin flow to generate motion. Many cell types, including cancer cells, can assemble a cortical flow engine to migrate under confinement and low adhesion *in vitro*, but it remains unclear whether this engine is endogenously utilized *in vivo.* Moreover, in the context of a changing environment, it is not known how upstream regulation can set in motion and sustain a mutual feedback between flow and polarity. Here, we establish that *Drosophila* primordial germ cells (PGCs) utilize cortical flow driven amoeboid migration and that flows are oriented by external cues during developmental homing *in vivo*. The molecular basis of flow modulation is a phosphoregulated feedback loop involving RhoGEF2, a microtubule plus-end tracking RhoA specific RhoGEF, enriched at the rear of PGCs. RhoGEF2 depletion slows and disorganizes cortical flow, reducing migration speed, while RhoGEF2 activation accelerates cortical flow, thereby augmenting myosin II polarity and migration speed. Both perturbations impair PGC pathfinding, suggesting cortical flows must be tuned for accurate guidance. We surprisingly find that RhoGEF2 polarity and activation are independent of upstream canonical Gα_12/13_ signaling. Instead, its PDZ domain and conserved RhoA binding residues in its PH domain are required to establish a positive feedback loop that augments its basal activity. Upstream regulation of this feedback loop occurs via AMPK dependent multisite phosphorylation near a conserved EB1 binding SxIP motif, which releases RhoGEF2 from EB1 dependent inhibition. Thus, we reveal cortical flows as versatile, tunable engines for directed amoeboid migration *in vivo*.

## Introduction

A shared mechanism amongst most migrating cells is the coupling of retrograde actin flow to the environment to generate forward motion. Retrograde actin flow is generally confined to the leading lamellipodia and lamellum in slower, adhesion dependent mesenchymal cells^1^, while in a subset of rapid amoeboid cells, cortical actin can flow across the entire cell cortex. This cell-scale cortical actin flow driven migration does not require canonical branched actin polymerization^2, 3^; instead it relies upon high levels of actomyosin contractility and low adhesion, and has been observed in a wide variety of cell types in 3D environments and/or under confinement, including breast cancer cells^3^, zebrafish progenitor cells^4, 5^, melanoma cells^6^, carcinosarcoma cells^2^, and various mesenchymal, immune, and epithelial cell lines^7–9^. Whether cells endogenously utilize this migration mode *in vivo* remains less certain, as previous, definitive observations in zebrafish embryos required ectopic induction of contractility via expression of constitutively active RhoA or wounding^4, 10^. Nevertheless, given the ubiquitous nature of this migration mode, its morphological hallmarks in migrating cancer cells in live mice^11^, and its rapid induction by purely environment factors^7^, understanding its molecular underpinnings and regulation will shed light on one of the most versatile means to migrate.

An intriguing question is how cortical flows are maintained and organized in migrating cells on a minutes to hours time scale. Cortical actin flows are drawn towards regions of high actomyosin contractility^12^ and are thought to be stabilized by the advection of front-back polarity factors, such as myosin II^4, 7, 13, 14^, as well as F-actin polymerization at the front and depolymerization at the rear^3, 15^. Upstream regulation of cortical flow, however, remains elusive, but is likely to depend on the conserved small Rho GTPase RhoA, as an LPA -> RhoA axis was required for its induction^4^ and pharmacological inhibition of its canonical downstream targets Dia^3^ and ROCK^3, 4, 7^ perturbs this migration mode. Current studies suggest external cues, such as confinement, induce but do not orient cortical flow driven amoeboid migration as a means to escape crowded envrionments^4, 5, 7, 9^. However, external guidance cannot be definitively ruled out, as the ectopic perturbations necessary to induce this migration mode thus far may override sensitivity to endogenous cues.

Primordial germ cells (PGCs) in many species are tasked with migrating across a complex, crowded cellular landscape that continuously evolves as development proceeds^16, 17^. As such, they require a flexible migration strategy that enables rapid movement through different cells expressing divergent cell adhesion molecules. One such strategy has been outlined in zebrafish PGCs, where F-actin accumulation templates the site of bleb formation and the subsequent local retrograde cortical actin flow following bleb expansion generates the necessary friction for forward movement^10, 18, 19^. Different strategies are likely to exist for other PGCs, including those in *Drosophila,* as in contrast to zebrafish PGCs, the activity of the small Rho GTPase Rac1 is not polarized^20^ and expression of a dominant negative Rac1 does not impair trans-epithelial PGC migration^21^.

In this study, we establish that Drosophila PGCs utilize a cortical flow engine to migrate during guided developmental homing *in vivo* and outline an upstream molecular pathway that modulates cortical flow that is necessary for accurate guidance.

## Results

### PGCs utilize retrograde cortical actin flows to directionally migrate *in vivo*

*Drosophila* PGCs maintain a spherical morphology while rapidly migrating through diverse adhesive cellular substrates during development. This migratory morphology is characteristic of a subtype of amoeboid migration involving rear contractility directed cortical actin flow employed by epithelial cells^8^ and cancer cells^3^ in 3D matrix, zebrafish progenitor cells upon wounding in *vivo*^4^, and a variety of cell lines upon confinement and low adhesion^7^. To determine whether cortical flows are similarly utilized by PGCs to migrate *in vivo*, we overexpressed the F-actin binding protein utrophin-GFP in embryos and imaged its dynamics along the dorsal cortex of PGCs, identified by a PGC specific membrane marker (tdKatushka2-CAAX), during directed migration toward the lateral mesoderm. With a fine temporal resolution (2.5 sec every frame), we observed a striking retrograde flow of cortical actin clusters commensurate with PGC movement in the opposite direction (Fig.1a, Supplementary Movie 1). Cortical actin flow speeds were consistently faster than PGC migration speeds (Fig. 1b), suggesting that these flows were slipping. Cortical actin flows can advect actin binding proteins, such as myosin II, toward the cell rear to establish and maintain front-rear migratory polarity^4, 7, 13, 14^. To determine whether myosin II underwent a similar retrograde flow as actin during PGC migration, we imaged PGCs expressing a myosin II 3x-GFP transgene under endogenous regulation. Although we were unable to resolve myosin II clusters on the dorsal surface of PGCs, we observed the retrograde flow of myosin II in the medial cortex near the cell rear during migration (Fig. 1c, Supplementary Movie 2). As opposed to actin flow, myosin II flow speed was on par with migration speed (Fig. 1d), which likely reflects the reduced cortical flow speeds near the cell rear, as observed in other amoeboid migrating cells^4, 7^, where our measurements were taken. We conclude that WT PGCs utilize cell scale retrograde cortical actin and myosin II flow to directionally migration *in vivo*, suggesting this migration mode can be oriented by external cues.

**Fig. 1.**
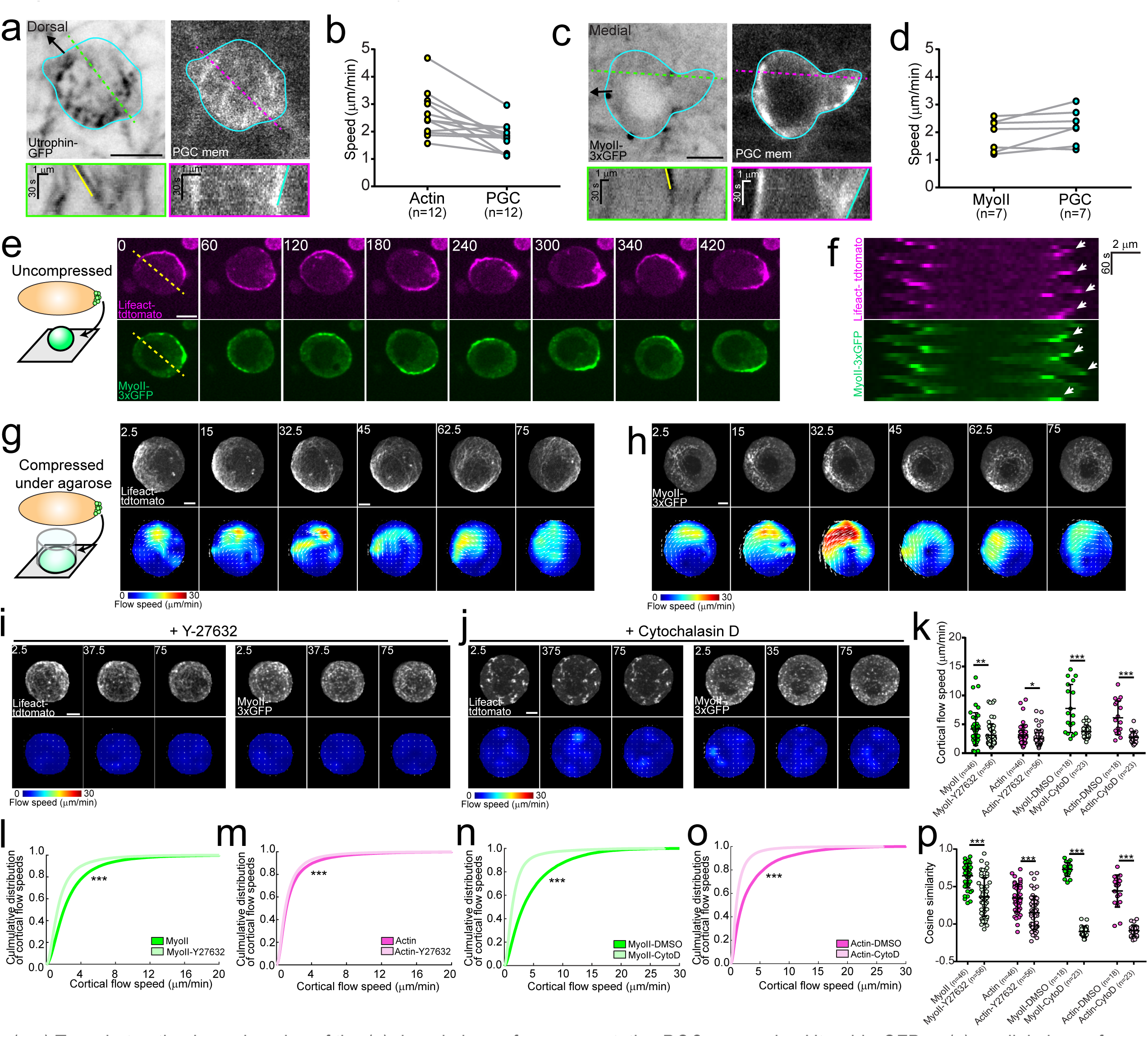
PGC cortical flow dynamics. (**a,c**) Two photon timelapse imaging of the (**a**) dorsal plane of a representative PGC expressing Utrophin-GFP or (**c**) medial plane of a representative PGC expressing myoII-3xGFP while co-expressing tdKatushka2-CAAX (PGC specific membrane marker). Cyan outline traces the boundary of the PGC as defined in the membrane image. Green and magenta dashed lines indicate where kymographs below the respective images were taken. The yellow line in the kymograph indicates retrograde flow of an actin cluster (**a**) or myosin II foci (**c**) while the cyan line tracks the rear of the cell. Scale bars, 5 μm. (**b,d**) Quantification of actin flow vs cell speeds (**b**) or myosin II flow vs cell speeds (**d**). The number of cells analyzed are indicated. (**e**) Timelapse imaging of an extracted PGC on coverglass express- ing lifeact-tdtomato and myoII-3xGFP. The yellow dashed line indicates where the kymograph in (**f**) was taken. (**f**) Kymograph from (**e**) with white arrows indicating periodic enriched F-actin and myosin II as flows sweep around the PGC. (**g-h**) Top panels- Timelapse imaging of a representative extracted PGC expressing lifeact-tdtomato (**h**) and myoII-3xGFP (**g**) under agarose. Bottom panel- PIV flow analysis between the image above and the previous time point. Flow speed is color coded with the indicated color bar. White arrows are the flow vectors at the indicated positions scaled to flow magnitude. (**i-j**) Top panels- Timelapse imaging of a representative extract- ed PGC expressing lifeact-tdTomato (left set) and MyoII-3xGFP (right set) under agarose with Y-27632 (**i**) or Cytochalasin D (**j**). Bottom panel- PIV flow analysis between the image above and the previous time point. Flow speed is color coded with the indicated color bar. White arrows are the flow vectors at the indicated positions scaled to flow magnitude. (**k**) Quantification of mean cortical flow speed of myosin II and actin under the indicated treatment conditions. The number of cells analyzed in each condition is indicated. Error bars are S.D. (**l-o**) Culmulative distribution of flow speeds of myosin II (**l,n**) and actin (**n,o**) after indicated drug treatment. (**p**) Quantification of mean cosine similarity between myosin II and actin vectors over consecutive time points under the indicated treatment conditions. Error bars are S.D. Time is in seconds in all images. Scale bars (**e-p**), 10 μm.*=p<.05, **=p<.01, and *** = p<.001.

### Extracted PGCs maintain cortical flows dependent on actomyosin contractility and actin polymerization

Observing cortical actin flows in migrating cells *in vivo* remains technically challenging^4^ and cells deep within tissues are not accessible to pharmacological perturbations. Thus, we developed methods to rapidly extract PGCs from stage 5 embryos to observe PGC dynamics *in vitro* (Fig. 1e-f, Fig. S1a-f, See materials and methods). Extracted PGCs expressing lifeact-tdTomato and myosin II-3xGFP plated onto uncoated glass in serum-free medium maintained their spherical morphology and exhibited 4 classes of cell behaviors- (1) striking periodic, circular flows of cortical actin and myosin II (Fig. 1e-f, Supplementary Movie 3) with a mean period of 86 +/- 21.7 seconds (S.D.) (41%, 78/188 cells), (2) stochastic accumulations of myosin II along the cortex (29%, 54/184 cells) (Fig. S1a-b), (3) blebbing (19%, 35/184) (Fig. S1c-d), and (4) inactive, where myosin II remained cytoplasmic (11%, 21/184) (Fig. S1e-f). Thus, as opposed to other cells which require an ectopic increase in contractility and/or confined environment to observe cortical flows^3, 4, 6–8^, the majority of extracted PGCs inherently exhibit continuous cell-scale cortical flow.

Placing PGCs under agarose substantially increased the surface area of the ventral cortex (nearest to glass), allowing us to image the flow of actin and myosin II cortical networks under high spatiotemporal resolution (Fig. 1g-h, Supplementary Movie 4). Particle image velocimetry (PIV) analysis indicated that myosin II foci traveled with the cortical actin network in sweeping, circular flows across the cell, mirroring our observations in uncompressed PGCs (Fig. 1g-h). Cortical actin and myosin II flow speeds reached upwards of 30 μm/min (Fig. 1g-h), an order of magnitude greater than the mean flow speeds we measured *in vivo*. These higher speeds are likely due to an uncoupling between actin flow and the environment, as PGCs remained stationary in these conditions, and may also arise from increased contractility under confinement, as observed in a variety of other cells^5, 7^.

Our ability to extract PGCs for *in vitro* experimentation further allowed us to determine the molecular requirements for cortical flow. Cortical flows have previously been shown to be dependent upon cortical contractility and actin polymerization^3, 4, 7^. Reducing actomyosin contractility via inhibition of the canonical upstream activator of myosin II, ROCK, with a strong dose of Y-27632 (100 μM) lead to a significant reduction in mean cortical actin and myosin II flow speeds, a significant shift in the cumulative distribution of all flow speeds, and a decrease in flow coordination (assessed by cosine similarity between the vectors at the same XY coordinate over consecutive time points) in the bulk population (Fig. 1i, k-m, p, Supplementary Movie 5). However, we did observe cells which appeared to have WT flow speeds and organized flows (Fig. 1k,p), suggesting PGCs may utilize other kinases to activate myosin II or possess alternative strategies to generate flow. In contrast to the heterogeneity we observed with ROCK inhibition, inhibiting actin polymerization with cytochalasin D fractured the actin cortex and lead to diffusive myosin II movements, resulting in a significant decrease in the mean cortical flow speed, a shift in the cumulative distribution of flow speeds, and disorganized flow in all cells as compared to control (Fig.1 j-k,n-p, Fig. S1g, Supplementary Movie 6-7). Taken together, these results suggest extracted PGCs maintain intrinsic cortical actin and myosin II flows observed *in vivo* which are dependent upon contractility and actin polymerization.

### RhoGEF2 is enriched at the rear of PGCs throughout developmental migration

Given that PGCs are guided by external cues during development^16^, there must be an upstream module to control cortical flow in PGCs. In migrating amoeboid cells, actin flow speed can be tuned by G protein-coupled receptor (GPCR) signaling but the downstream pathways connecting the two remain unclear^22^. One pathway likely involved is the conserved contractility pathway downstream of the small Rho GTPase, RhoA, whereby GTP-bound, active RhoA binds and relieves auto-inhibition of ROCK, which then phosphorylates and activates myosin II. Cortical flow is drawn towards regions of high actomyosin contractility^12^ and we have previously shown that RhoA is active at the rear of migrating PGCs^20^. We thus hypothesized that regulation of a RhoA activator, a Rho guanine nucleotide exchange factor (GEF), may allow cortical flow modulation.

To identify candidate RhoGEFs involved in PGC migration, we assessed available GFP-tagged RhoGEFs via live imaging for posterior enrichment. During stage 9 of embryogenesis, PGCs reside in a cluster and radially align front-rear polarity outwards under the guidance of the GPCR, Tre1, thus providing a robust readout for posterior proteins which enrich in the center of the cluster^20^. We discovered that sfGFP tagged RhoGEF2, a RhoA specific RhoGEF, driven by a ubiquitous *squash* (*Drosophila* myosin II regulatory light chain) promoter was concentrated at the rear of polarized PGCs (Fig. 2a,c). This enrichment was not due to overexpression, as a sfGFP-RhoGEF2 fosmid, where RhoGEF2 is driven by endogenous regulation, had a similar enrichment and immunostaining of endogenous RhoGEF2 revealed a similar posterior polarization in migrating PGCs (Fig. S2). In the absence of Tre1, PGCs are unable to separate^20, 21, 23–25^ and exhibit extensive intracluster motility and cell turning. As a result, rear enriched proteins, such as myosin II, now appear at the edge of the cluster. We similarly observed that polarized RhoGEF2 appeared at the periphery of clusters in Tre1 mutants, although it was not as strongly polarized as in WT PGCs (Fig. 2b-d). This suggests RhoGEF2 is involved in the basal motility of PGCs and may be further recruited under GPCR guidance. Two-photon live imaging of migrating PGCs during various stages of development further confirmed that RhoGEF2 was dynamically enriched at the rear during all phases of PGC migration (Fig. 2e-f, Supplementary Movie 8-9).

**Fig. 2.**
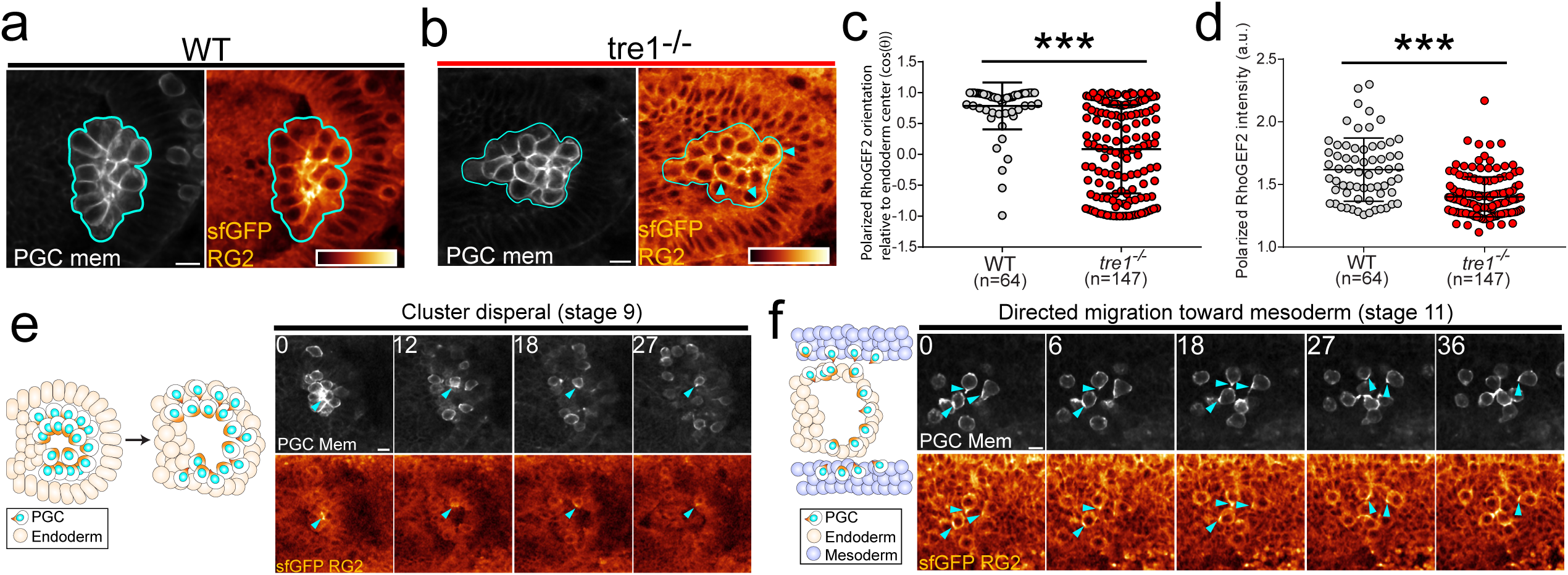
RhoGEF2 is posteriorly enriched throughout developmental migration. (**a-b**) Representative two photon image of the central plane of a WT (**a**) or tre1^-/-^ (**b**) PGC cluster expressing sfGFP-RhoGEF2 (RG2) and tdKatushka2-CAAX (PGC membrane marker). Cyan outlines the PGC cluster while cyan arrows indicate regions of polarized RhoGEF2. sfGFP RhoGEF2 is color coded with the indicated color bar. (**c,d**) Quantification of polarized RhoGEF2 orientation relative to the center of the cluster (**c**) or Polarized RhoGEF2 intensity (**d**). The number of PGCs analyzed are indicated from n=7 embryos (WT) and n=15 embryos (tre1^-/-^). Error bars are S.D. (**e-f**) Two photon timelapse imaging of representative PGCs expressing sfGFP-RhoGEF2 and tdkatushka2-CAAX during cluster dispersal (n = 7 embryos imaged) (**e**) and migration toward mesoderm (n=6 embryos imaged) (**f**). Cyan arrows indicate regions of polarized RhoGEF2 accumulation. sfGFP-RhoGEF2 is color coded with the color bar in **a**.Time is in minutes in all images. All scale bars, 10 μm.*** = p<.001.

### RhoGEF2 regulates cortical flow and is necessary for accurate guidance

To assess whether RhoGEF2 is necessary for PGC migration, we utilized a GFP degradation system, degradFP^26^, to degrade maternal sfGFP-RhoGEF2 in a genetic background where sfGFP-RhoGEF2 is the sole source of RhoGEF2 in the embryo. We created a PGC targeted lexA inducible degradFP transgene and drove it with a newly constructed maternal LexA driver (Maternal tubulin promoter). As RhoGEF2 is necessary for somatic cellularization^27^, we rescued any somatic defects via overexpression of an untagged RhoGEF2 using an early soma specific Gal4 driver, *nullo*-Gal4. The use of orthogonal LexA and Gal4 systems prevented any potential crosstalk between these constructs. Live imaging indicated that RhoGEF2 was significantly depleted (reduced to ∼35%) prior to the dispersal of PGC clusters, allowing us to assess its role in directed migration dependent cluster dispersal and subsequent steps in developmental migration (Fig. S3). RhoGEF2 depletion significantly deceased the rate at which PGCs detached from clusters and transmigrated through the endoderm (Fig. 3a-c, Supplementary Movie 10), suggesting defects in generating and orienting motility. Following delayed cluster detachment, we similarly observed that PGC migration speed was reduced during directed migration towards the mesoderm (Fig. 3d-f, Supplementary Movie 11). Consequently, the number of PGCs which failed to arrive at the gonad at stage 14, when developmental migration has concluded, increased (Fig. 3g-i).

**Fig. 3.**
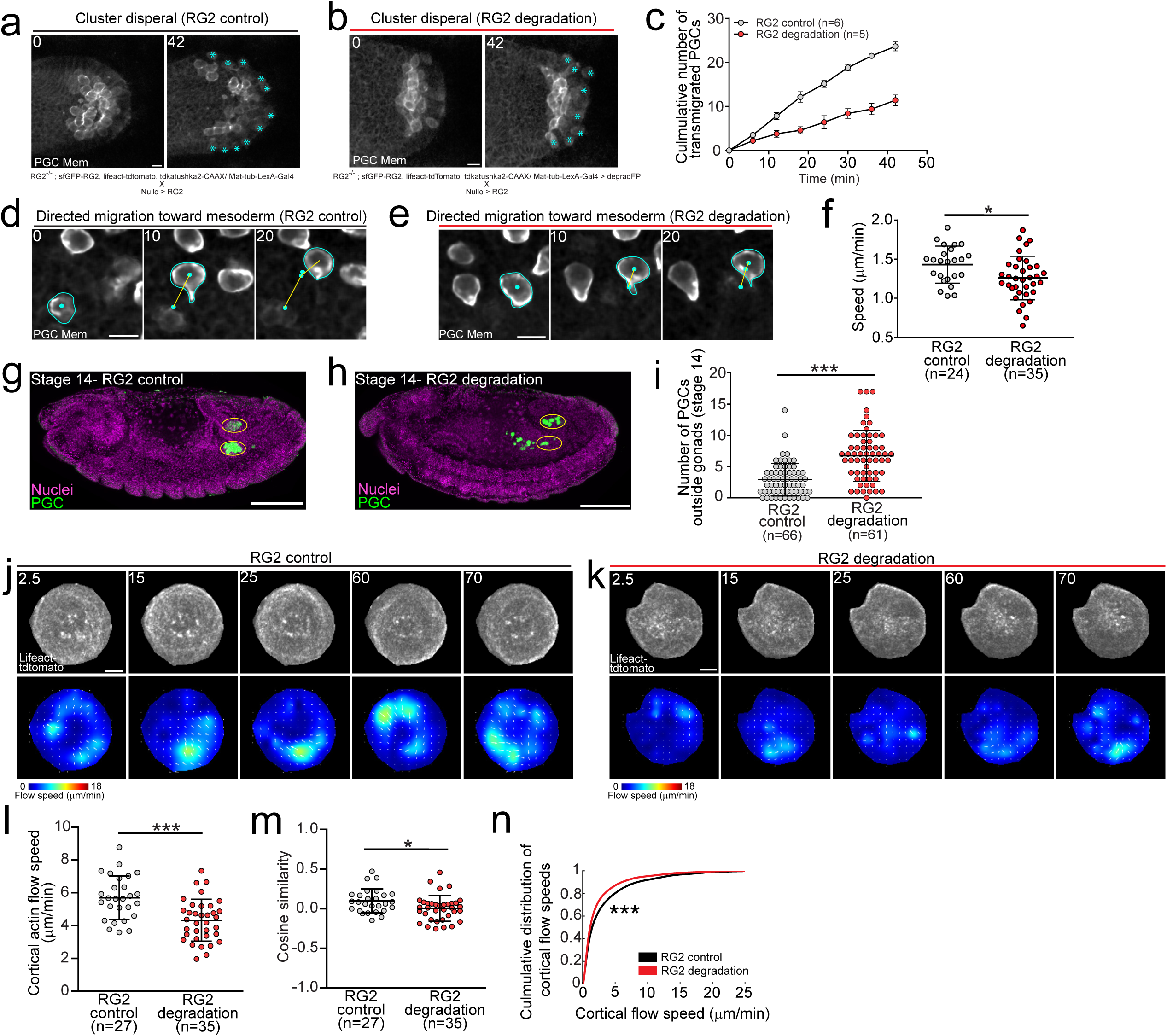
RhoGEF2 regulates cortical flow and is required for accurate guidance. (**a-b,d-e**) Representative two photon timelapse imaging of PGCs expressing lifeact-tdtomato (not shown) and tdkatushka-CAAX (membrane marker) under the indicated conditions during cluster dispersal (**a-b**) or directed migration toward the mesoderm (**d-e**). Genotypes are listed below images. Cyan asterisks mark transmigrated PGCS in **a-b**. Cyan outlines cell membranes while cyan dots track PGC nuclei in **d-e**. (**d-e**) Yellow lines are cell tracks between indicated times. Scale bars, 10 μm. Times are in minutes. (**c**) Quanti- fication of the total number of transmigrated PGCs in the indicated conditions over time. The number of embryos analyzed is indicated. Error bars are SEM. (**f**) Quantification of cell speed under the indicated conditions. The number of PGCs analyzed is indicated from n=6 embryos (RG2 control) and n=5 embryos (RG2 degradation). Error bars are S.D. (**g-h**) Representative immunofluorescence images of stage 14 embryos under the indicated conditions. Yellow ovals indicate the gonads. Scale bars, 100 μm. (**i**) Quantification of the number of PGCs outside gonads in indicated conditions. The number of embryos analyzed is indicated. Scale bars are S.D. (**j-k**) Top panel- timelapse imaging of representative extracted PGCs under agarose with the indicated experimental conditions expressing lifeact-tdTomato. Bottom panel- PIV flow analysis between the image above and the previous time point. Flow speed is color coded with the indicated color bar. White arrows are the flow vectors at the indicated positions scaled to flow magnitude. Scale bars, 5 μm. (**l-m**) Quantification of mean cortical actin flow speed (**l**) and mean cosine similarity between vectors over consecutive time points (**m**) in the indicated conditions. The number of cells analyzed is indicated. Error bars are S.D. (**n**) Culmulative distributions of flow speeds in the indicated conditions. *=p<.05 and ***=p<.001.

We hypothesized that the migration defects we observed after RhoGEF2 depletion were due to perturbed cortical flow, as a presumed flatter actomyosin contractility gradient in these PGCs could less efficiently drive and organize flow. To test this idea, we extracted control and RhoGEF2 depleted PGCs from embryos and imaged cortical actin flow using our agarose assay. Cortical flow speeds were significantly slower and more disorganized in RhoGEF2 depleted PGCs, suggesting RhoGEF2 plays an important role in organizing and driving flow (Fig. 3j-n, Supplementary Movie 12). In sum, we find that RhoGEF2 dependent cortical flow modulation is necessary for accurate PGC migration *in vivo*.

### Excess RhoGEF2 activation enhances cortical flow and polarity but impairs guidance

We next asked whether enhancing RhoGEF2 activity would impair migration. RhoGEF2 is a microtubule plus-end tracking RGS-homology (RH) RhoGEF and is best known for its role in gastrulation, where it lies downstream of a well described linear cascade (Fig. 4a). At the top of this pathway, the ligand, Fog, activates the GPCR, Mist, leading to Gα_12/13_ (known as concertina in *Drosophila*) activation, which then binds and activates RhoGEF2^28–32^. Given that RhoGEF2 is the only known target of Gα_12/13_ in *Drosophila*, expression of a constitutively active Gα_12/13_ (Gα_12/13-_Q303L) would selectively active RhoGEF2^33^. Thus, we created a PGC targeted Gα_12/13-_Q303L transgene, drove it in the embryo maternally with a maternal tubulin Gal4-VP16, and examined its effects on RhoA activity with live two-photon imaging of a RhoA activity sensor (Anillin-RhoA binding domain (RBD) fused to tdTomato). Contrary to our expectation of global activation, upon examining stage 9 PGC clusters, we observed increased RhoA activation solely in the cluster center, suggesting that global RhoGEF2 activation did not disrupt but rather enhanced front-back polarity (Fig. S4). This enhanced posterior RhoA activity lead to a concomitant increase in myosin II enrichment in the cluster center, as assessed with a transgenic line expressing myosin II-tdTomato and tdkatushka2-CAAX (membrane marker) in PGCs (Fig. 4b-d). Notably, live imaging of PGC migration toward the mesoderm further confirmed this enhanced myosin II polarity in individual cells and indicated a significant increase in migration speed (Fig. 4e-g, Supplementary Movie 13). This enhanced front-back polarity and speed, however, severely disrupted PGC guidance to the gonad (Fig. 4h-i), suggesting potential defects in orienting or stopping migration.

**Fig. 4.**
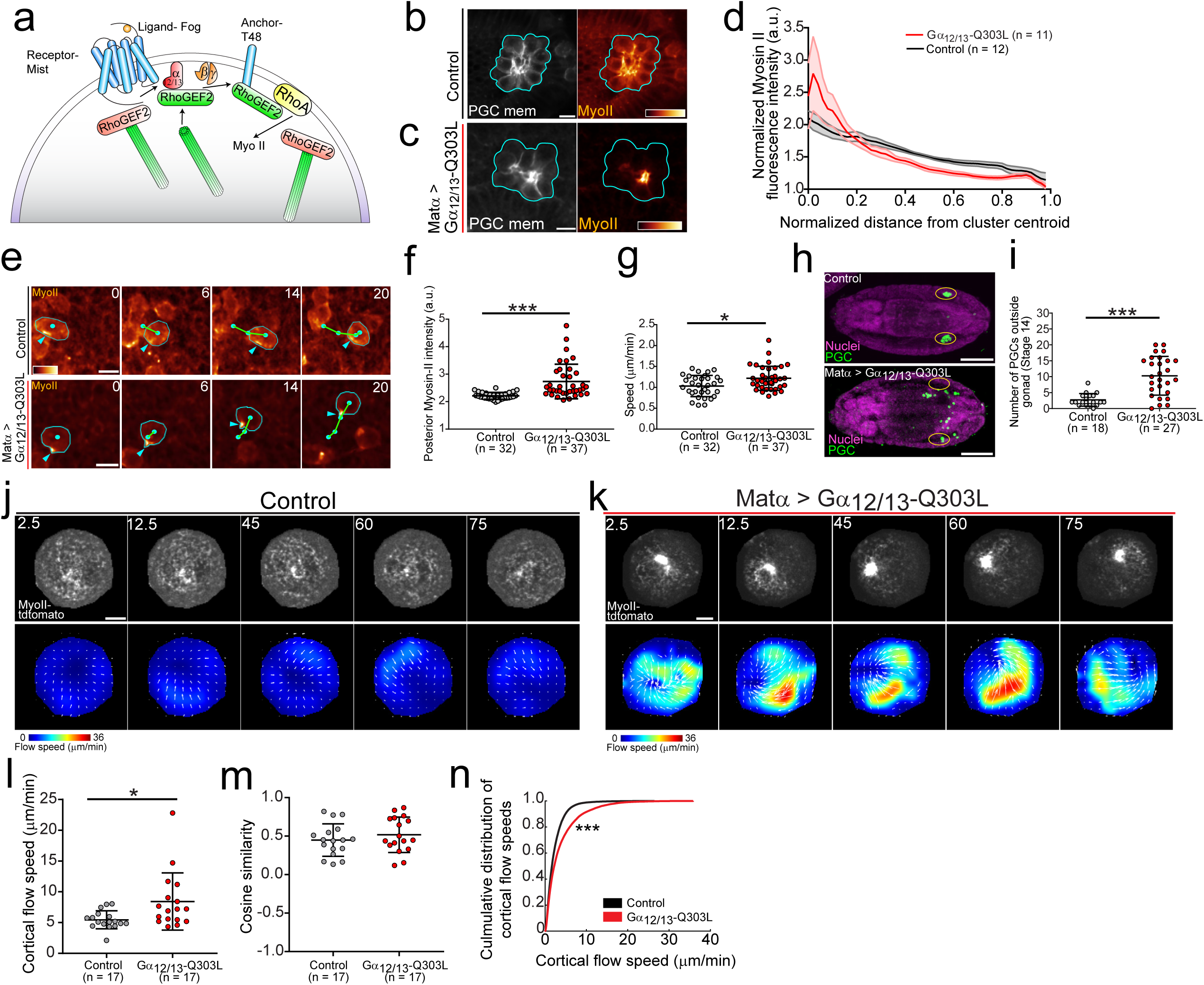
Excess RhoGEF2 activati on enhances polarity and speed but impairs homing. (a) Model of the canonical RhoGEF2 pathway in gastrulation. (**b-c**) Representative two photon image of a central plane from PGC clusters expressing myosin II-tdtomato and tdKatushka2-CAAX under the indicated conditions. Cyan outlines the perimeter of the cluster and myosin II-tdtomato is pseudocolored with the indicated color bar. Scale bar, 10 μm. (**d**) Quantification of myosin-II tdtomato intensity as a function of distance from the cluster centroid. The number of embryos analyzed is indicated. Error bars are SEM. (**e**) Representative two photon timelapse imaging of PGCs expressing myosin II-tdtomato migrating to the mesoderm in the indicated condtions. Cyan outlines the PGC periphery, cyan dots track PGC position, and cyan arrows indicate regions of polarized myosin II. Green lines are the cell tracks between the indicated time periods. Times are in minutes. Scale bars, 10 μm. (**f-g**) Quantification of polarized myosin II intensity (**f**) and speed (**g**) in the indicated conditions. The number of cells analyzed is indicated from n=4 embryos (control) and n=5 embryos (Gα12/13-Q303L overexpression). Error bars are S.D. (**h**) Representative immunofluorescence images of stage 14 embryos from the indicated conditions. Yellow ovals mark the gonads. Scale bars, 100 μm. (**i**) Quantification of the number of PGCs outside gonads in indicated conditions. The number of embryos analyzed is indicated. Error bars are S.D. (**j-k**) Top panel- Timelapse imaging of representative extracted PGCs under agarose with indicated experimental conditions expressing myosin II-tdTo- mato. Bottom panel- PIV flow analysis between the image above and the previous time point. Flow speed is color coded with the indicated color bar. White arrows are the flow vectors at the indicated positions scaled to flow magnitude. Scale bars, 5 μm. (**l-m**) Quantification of mean cortical actin flow speed (**l**) and mean cosine similarity between vectors over consecutive time points (**m**) in the indicated conditions. The number of cells analyzed is indicated. Error bars are S.D. (**n**) Culmulative distributions of flow speeds in the indicated conditions. *=p<.05 and ***=p<.001.

The enhanced myosin II accumulation we observed after Gα_12/13-_Q303L overexpression could arise from the acceleration of cortical flow, as faster cortical flows can steepen front-back gradients of actin binding polarity factors, such as myosin II, as they travel further for a given time period^13^. We tested this idea by extracting control and Gα_12/13-_Q303L overexpressing PGCs and imaging cortical myosin II flow. Indeed, activating RhoGEF2 lead to significant increase in cortical myosin II flow speeds without affecting flow organization and we identified large myosin II foci in Gα_12/13-_Q303L overexpressing PGCs (Fig. 4j-n, Supplementary Movie 14), in accord with our observations *in vivo* (Fig. 4e-f). Our results collectively indicate that RhoGEF2 activity levels scale cortical flow and migration speeds and further suggest cortical flow speeds must be tuned for accurate homing.

### RhoGEF PDZ and PH domains are required for polarity and migration

We next sought to determine how RhoGEF2 is polarized and activated in migrating PGCs (Fig. 5a). Like many RhoGEFs, RhoGEF2 is a large multi-domain protein and possesses four notable domains- an N terminal PDZ domain, a RGS domain responsible for binding Gα_12/13_, a C1 domain, and a tandem DH-PH domain necessary for catalytic activity (Fig. 5b). Previous work has shown that the PDZ domain regulates RhoGEF2 localization through interaction with transmembrane protein anchors, such as T48 during gastrulation^34^ (Fig. 4a) and slam during cellularization^35^. However, these anchors are not expressed in PGCs^34, 36^. Moreover, although Gα_12/13_ is the canonical RhoGEF2 activator in *Drosophila*^31^, we have previously shown that Gα_12/13_ is not required for PGC migration^37^. Thus, an alternative localization and activation mechanism must exist in PGCs (Fig. 5a).

**Fig. 5.**
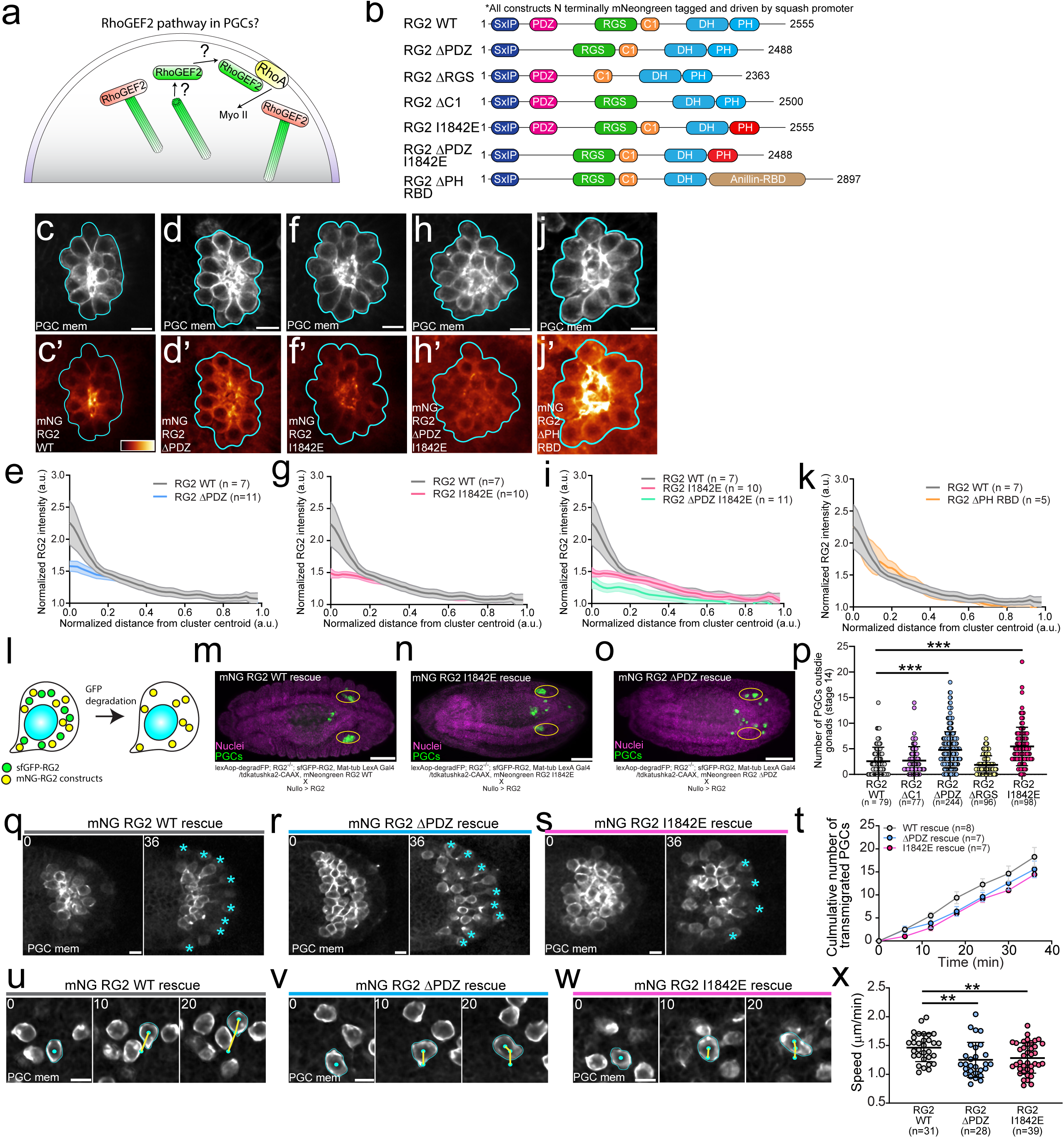
RhoGEF2 PDZ and PH domains are necessary for polarity and migration. (**a**) Model of the currently unknown RhoGEF2 pathway in PGCs. (**b**) Schematic of RhoGEF2 protein domains and the series of constructs created based on domain truncations and mutations. All constructs are driven by an ubiquitous *squash* (myosin II regulatory light chain) promoter and are N terminally tagged with mNeongreen. (**c-d,f,h,j**) Representative two photon image of a central plane from PGC clusters expressing the indicated mNeongreen tagged RhoGEF2 transgene and tdKatushka2-CAAX (PGC specific mem- brane marker). mNeongreen-RhoGEF2 transgenes are pseudocolored with the color bar in c. Cyan outlines the perimeter of the cluster. Scale bar, 10 μm. (**e,g,i,k**) Quantification of the indicated mNeongreen RhoGEF2 transgene intensity as a function of distance from the cluster centroid vs WT. The number of embryos analyzed is indicated. Error bars are SEM. (**l**) Schematic of the experimental scheme to degrade sfGFP-RhoGEF2 and leave PGCs with a given mNeongreen-RhoGEF2 transgene. (**m-o**) Representative immuno- fluorescence images of stage 14 embryos under the indicated rescue conditions. Genotypes are stated below. Yellow ovals mark the gonads. Scale bars, 100 μm. (**p**) Quantification of the number of PGCs outside gonads in indicated rescue conditions. The number of embryos analyzed is indicated. Error bars are S.D. (**q-s**) Two photon timelapse imaging of representative PGC clusters expressing tdKatushka2-CAAX (PGC membrane marker) dispersing under the indicated rescue conditions. Cyan asterisks mark transmigrated PGCs. (**t**) Quantification of the total number of transmigrated PGCs under the indicated rescue conditions over time. The number of embryos analyzed is noted. Error bars are SEM. (**u-w**) Two photon timelapse imaging of representative PGCs migrating towards the mesoderm under the indicated rescue conditions. Cyan outlines cell membranes while cyan dots track PGC position. Yellow lines are cell tracks between indicated times. Scale bars, 10 μm. Times are in minutes. (**x**) Quantification of mean cell migration speed under the indicated rescue conditions. The number of PGCs analyzed is noted from n=6 embryos (RG2 WT), n=4 embryos (RG2 ΔPDZ), and n=7

To identify which domains are necessary for RhoGEF2 function in PGCs, we created a panel of transgenic lines harboring ubiquitous *squash* promoter driven mNeongreen tagged RhoGEF2 lacking the PDZ, RGS, or C1 domain (Fig. 5b). In insect S2 cells, all constructs retained their ability to track microtubule plus-ends and activated myosin II in a similar dose dependent manner, with the highest activation levels with the ΔRGS construct, suggesting these truncations did not grossly affect RhoGEF2 function (Fig. S5a-c). The PDZ and RGS domains were, however, essential for RhoGEF2 function *in vivo*, while the ΔC1 transgene provided a comparable rescue to WT (Table S1). These differences did not arise from expression levels, as all transgenes were expressed at comparable levels in embryos (Fig. S5d-e). Live imaging in stage 5 embryos also confirmed that only the ΔPDZ transgene was not enriched on cellularization furrows (Fig. S5f-g), in line with previous work indicating this is a PDZ domain dependent process^35^. Subsequent live imaging of stage 9 PGC clusters expressing these RhoGEF2 transgenes and a PGC specific membrane marker (tdKatushka2-CAAX) revealed that only the PDZ domain appreciably contributed to RhoGEF2 polarization (Fig. 5c-e, Fig. S5h-k), suggesting a congruence with other developmental processes, albeit with a different putative transmembrane anchor.

Given that RhoGEF2 still polarized in PGCS without the PDZ domain, we next asked what other mechanisms could regulate localized RhoGEF2 function. Recent work has shown that the PH domain of most RH-RhoGEFs can bind active, RhoA-GTP *in vitro* at a distal site from the catalytic DH domain, providing a localization based positive feedback^38^. Dampening this feedback by mutating conserved hydrophobic residues critical for this interaction in the PH domain substantially reduces RH-RhoGEF overexpression induced RhoA activation in mammalian cells. Interestingly, for PDZ-RhoGEF, this feedback loop can potentiate its basal activity up to 40 fold in the absence of Gα_12/13_^38^, which regulates its localization but does not enhance its exchange activity^39^. An analogous mechanism could explain how RhoGEF2 augments its basal activity to regulate PGC migration without requiring Gα_12/13_.

To assess whether this feedback loop exists for RhoGEF2, we first asked whether the RhoGEF2 PH domain could bind RhoA-GTP in an *in vitro* binding assay. An *in vitro* translated RhoGEF2 PH domain bound purified GST-RhoA-GDP and GST-RhoA-GTPγS but not GST, suggesting this interaction was specific for RhoA (Fig. S6a-c). Introducing a point mutation (I1842E) in one of two conserved hydrophobic residues in the PH domain critical for RhoA-GTP binding in other RH-RhoGEFs decreased both affinities but had a stronger effect on RhoA-GTP binding, suggesting these residues are also important for RhoGEF2 PH-RhoA interaction. Overexpression of mNeongreen-RhoGEF2 with either PH point mutation (I1842E, F1840A) expectedly reduced myosin II activation in S2 cells at comparable expression levels vs. WT but did not affect microtubule plus-end tracking (Fig. S6d-f). The Glotzer group has also shown that mutating these residues suppresses the lethality associated with constitutive expression of an optogenetic RhoA activation system utilizing the DH-PH domain from RhoGEF2^40^.

To determine whether this feedback was functionally significant *in vivo*, we established *squash* promoter driven mNeongreen tagged RhoGEF2 transgenic lines containing these PH domain mutants (Fig. 5b). Both RhoGEF2 PH mutant lines were expressed at similar levels as WT but could not rescue RhoGEF2 mutants (Fig. S5d-e, Table S1), suggesting the RhoGEF2 PH-RhoA interaction is critical for RhoGEF2 function. These PH mutations did not perturb enrichment on cellularization furrows in stage 5 embryos (Fig. S6g-h). In stage 9 PGC clusters, both PH mutations reduced RhoGEF2 polarization to a similar extent as the ΔPDZ construct, suggesting this RhoA binding feedback contributes to RhoGEF2 localization and likely activation in PGCs (Fig. 5f-g, Fig. S6i-j). To assess whether the PDZ and PH domains localize RhoGEF2 in parallel, we generated a double mNeongreen ΔPDZ, I1842E RhoGEF2 transgenic line and observed a further reduction in RhoGEF2 polarity in PGC clusters than with either domain perturbation alone (Fig. 5b,h-i). Collectively, our results suggest RhoGEF2 is regulated in PGCs through two parallel mechanisms- (1) PDZ domain binding to an unknown anchor to enrich it at the cell rear and (2) PH domain binding to RhoA-GTP to amplify nascent sites of RhoA activity triggered by its basal activity.

If the PH domain acts to enhance RhoGEF2 activity by retaining it near its substrate, a domain with a similar function should be able to compensate for its loss. We tested this idea by generating a transgenic line where the RhoGEF2 PH domain was swapped with the RhoA-GTP binding domain (RBD) from Anillin (Fig. 5b). Although this transgene could not rescue lethality associated with a RhoGEF2 mutant, we interestingly found that it recapitulated WT levels of RhoGEF2 polarity in stage 9 PGC clusters (Fig. 5j-k), suggesting this is the main function of the PH domain.

We next sought to determine whether the same RhoGEF2 domains necessary for polarity were likewise important for PGC migration. Since the mNeongreen ΔPDZ, ΔRGS, and PH domain mutant RhoGEF2 transgenes could not rescue RhoGEF2 mutants (Table S1), we could not directly assess if these domains were critical for PGC migration without the confounding presence of the WT protein. To overcome this limitation, we rescued RhoGEF2 null embryos with sfGFP-RhoGEF2 while co-expressing a given mNeongreen RhoGEF2 transgene. We then removed sfGFP-RhoGEF2 with degradFP using the experimental scheme described earlier, allowing us to determine whether the remaining mNeongreen RhoGEF2 protein was sufficient for PGC migration (Fig. 5l). In accord with the polarity phenotypes we observed above, removing the PDZ domain or mutating the PH domain more severely disrupted PGC homing than any other domain perturbation (Fig. 5m-p). Live imaging further confirmed that these homing defects stemmed from a decreased cluster dispersal rate and slowed migration (Fig. 5q-x, Supplementary Movie S15-16). We propose that RhoGEF2 polarity and activity in PGCs require its PDZ and PH domains which directly influence migration prowess through cortical flow speed modulation.

### RhoGEF2-EB1 inhibition is relieved by phosphorylation

Our results thus far suggest RhoGEF2 activity must be finely tuned to achieve accurate migration, as increasing or decreasing its activity impairs PGC guidance. However, its known regulation involves Gα_12/13,_ which is not necessary for PGC migration (Fig. 4a). Interestingly, RhoGEF2 is inactive when bound to microtubule plus-ends and is liberated by Gα_12/13_, suggesting titration of free RhoGEF2 as a regulation strategy^41^ (Fig. 4a). RhoGEF2 possesses an N terminal SxIP motif (SKIP) that indirectly allows a variety of proteins to track microtubule plus-ends via interaction with EB1^42^, but it has not been determined whether this motif is required for RhoGEF2 plus-end tracking. To establish the functional significance of the SKIP motif in RhoGEF2, we introduced two point mutations previously shown to abrogate EB1 interaction (SKIP -> SKNN) (Fig. 6a). Live imaging of insect cells expressing mNeongreen RhoGEF2 SKNN and EB1-mScarlet revealed that this construct was unable to track plus-ends while the control construct did (Fig. 6b-c). These results suggest the RhoGEF2 SKIP motif mediates microtubule plus end tracking.

**Fig. 6.**
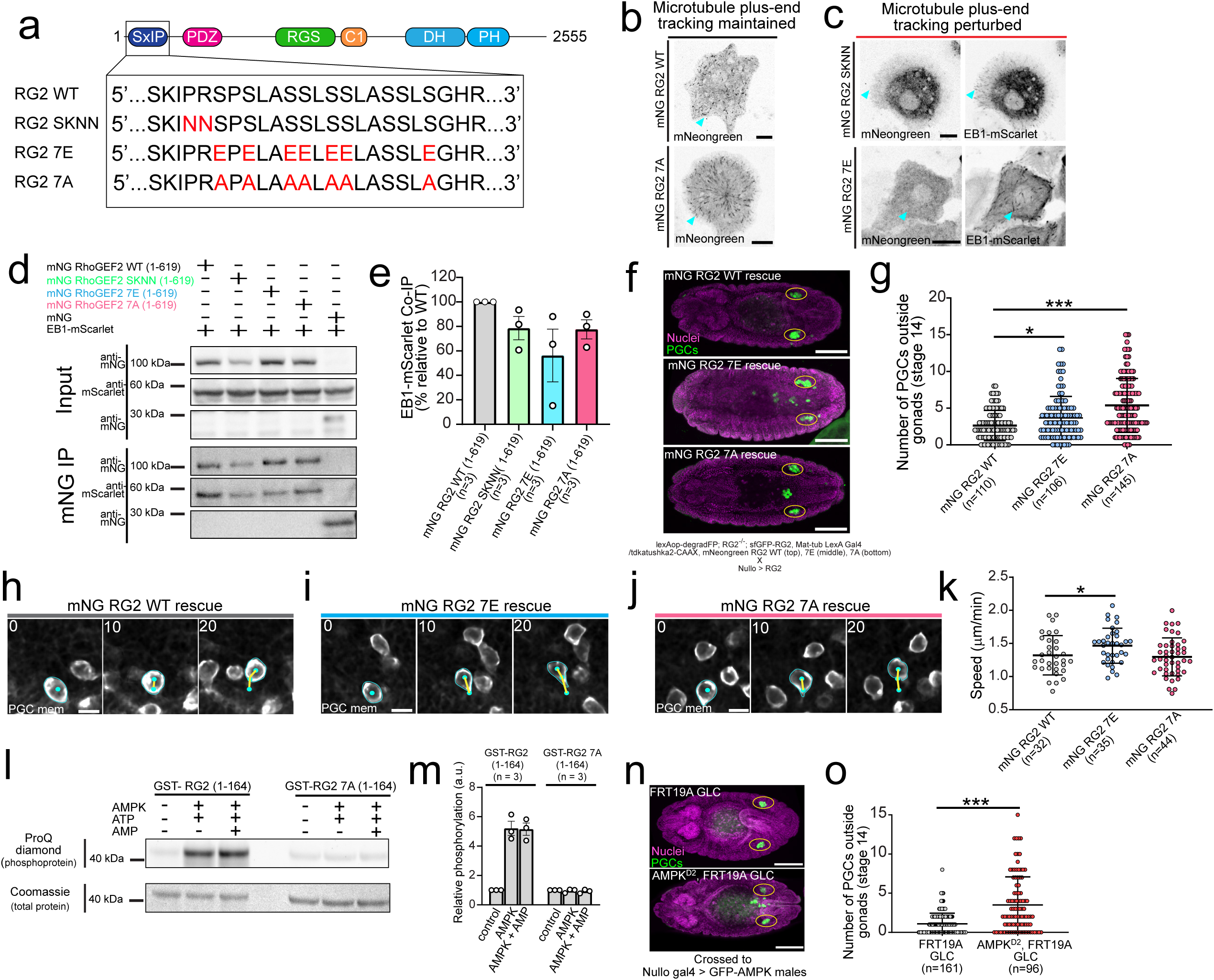
RhoGEF2 phosphoregulation titrates free RhoGEF2 for accurate migration. (a) Schematic of the RhoGEF2 SKIP motif and protein domains along with the series of constructs created with mutagenesis. All constructs are driven by a *squash* promoter and N terminally tagged with mNeongreen. (**b-c**) Representative image from S2 cells expressing the indicated mNeongreen RhoGEF2 constructs. EB1-mScarlet is co-expressed in (**c**) to identify microtubule plus-ends. Cyan arrows highlight microtubule plus-end tracking. Scale bars, 10 μm. (**d**) Representative immunoblot from a co-immunoprecipitation experiment from S2 cells overexpressing the indicated constructs. Cell lysates were immunoprecipitated with mNeongreen-trap and probed with the indicated antibodies. Immunoprecipitation, IP. (**e**) Quantification of EB1-mScarlet co-immunoprecipitation with the indicated mNeongreen RhoGEF2 constructs. Error bars are SEM. (**f**) Representative immunofluorescence images from stage 14 embryos under the indicated rescue conditions. Yellow ovals mark the gonads. Genotypes are listed below the images. Scale bars, 100 μm. (**g**) Quantification of the number of PGCs outside gonads in indicated conditions. The number of embryos analyzed is indicated. Error bars are S.D. (**h-j**) Two photon timelapse imaging of representative PGCs migrating towards the mesoderm under the indicated rescue conditions. Cyan outlines cell membranes while cyan dots track PGC position. Yellow lines are cell tracks between indicated times. Scale bars, 10 μm. Times are in minutes. (**k**) Quantification of mean migration speed under indicated rescue conditions. Error bars are S.D. The number of PGCs analyzed is noted from n = 5 embryos (mNG RG2 WT), n=5 embryos (mNG RG2 7E), and n=6 embryos (mNG RG2 7A). (**l**) Representative SDS-PAGE gel from an in vitro kinase assay with purified AMPK enzyme and GST-RG2 fusion proteins under the indicated conditions. The gel was initially probed with a ProQ diamond stain to visualize phoshorylated proteins and was subsequently stained with Coomassie blue to visualize total proteins. (**m**) Quantification of GST-RG2 protein phos- phorylation relative to control under the indicated conditions. Error bars are S.D. (**n**) Representative immunofluorescent images from stage 14 embryos with the indicated female genotypes. Yellow ovals mark the location of the gonads. Scale bars, 100 μm. GLC, germ line clone. (**o**) Quantification of the number of PGCs outside gonads in indicated conditions. The number of embryos analyzed is indicated. Error bars are S.D. *=p<.05, **=p<.01, and ***=p<.001.

We next asked how the RhoGEF2 SKIP motif is regulated. SxIP motifs are typically flanked by numerous serine residues, which, when phosphorylated, can sterically inhibit EB1 interaction, allowing upstream regulation by a kinase^42^. The RhoGEF2 SKIP motif is likewise surrounded by several serine residues, many of which are phosphorylated in the embryo, as indicated in a phospho-proteomic database from the Perrimon lab^43^ (Fig. 6a). To determine whether multi-site phosphorylation near the SKIP motif could impact RhoGEF2-EB1 interaction, we substituted glutamic acids or alanines for these seven proximal serines to create phosphomimetic (7E) and phosphonull (7A) RhoGEF2 constructs, respectively (Fig. 6a). Upon overexpression in insect S2 cells, all constructs, including RhoGEF2 SKNN, exhibited slightly higher levels of myosin II activation as compared to WT at intermediate levels, but this was only significant for RhoGEF2 SKNN. At high expression levels there were no significant differences with WT RhoGEF2, presumably because the buffering capacity of EB1 had been exceeded (Fig. S7a-b). Live imaging of insect cells co-expressing mNeongreen RhoGEF2 7E and EB1-mScarlet revealed that similar to RhoGEF SKNN, RhoGEF2 7E could not track microtubule plus ends (Fig. 6c), while the mNeongreen RhoGEF2 7A construct expectedly retained this ability (Fig. 6b). To further validate these conclusions, we performed co-immunoprecipitation experiments from S2 cells co-expressing truncated mNeongreen RhoGEF2 constructs with these mutations and EB1-mScarlet with mNeongreen itself as a control. These truncated proteins maintained the microtubule plus-end phenotypes observed with the full length protein (Fig. S7c-f). We found that the truncated RhoGEF2 SKNN, 7E, and 7A constructs all had a reduced affinity for EB1-mScarlet as compared to WT with the 7E construct exhibiting the lowest affinity overall (Fig. 6d-e). The reduced affinity of the 7A construct was unexpected, given its microtubule plus-end tracking in live imaging experiments (Fig. S7f), and may reflect a change in protein behavior when extracted from a cellular environment. Our results collectively suggest the free pool of signaling competent RhoGEF2 can be titrated by multi-site phosphorylation near the RhoGEF2 SKIP motif.

We next asked whether perturbing RhoGEF2-EB1 phosphoregulation had consequences for PGC migration. Following a similar strategy as above, we generated transgenic lines harboring mNeongreen RhoGEF2 SKNN, 7E, and 7A which expressed at comparable levels to the WT (Fig. S7g-h) and polarized to a similar extent as WT in PGC clusters (Fig. S7i-k). However, in contrast to the ΔPDZ, ΔRGS, PH domain mutant RhoGEF2 transgenes, these lines were able to rescue RhoGEF2 mutants, suggesting this phosphoregulation is not absolutely essential for RhoGEF2 function. Perturbing RhoGEF2-EB1 phosphoregulation, however, did impair PGC homing, as there were significantly more PGCs which did not arrive at the gonad in stage 14 embryos when we performed rescue experiments, suggesting this mechanism is necessary for PGC migration (Fig. 6f-g). To investigate why PGC guidance was disturbed, we performed two-photon live imaging of PGCs expressing RhoGEF2 7E and 7A during cluster dispersal and migration toward the mesoderm. Although the cluster dispersal rate was similar in all conditions (Fig. S7l-o), PGCs expressing RhoGEF2 7E migrated significantly faster than WT (Fig. 6h-k), suggesting the free levels of RhoGEF2 were tied to migration speed. The migration defects we observed in the RhoGEF 7A rescue experiments, which do not perturb PGC speed, could arise from defects in PGC migration in developmental steps we did not assess. Collectively, our findings suggest dynamic, phosphorylation-mediated control of free RhoGEF2 levels allows the dose dependent control of migration speed necessary for accurate migration.

### AMPK directly phosphorylates RhoGEF2 and is necessary for PGC guidance

Lastly, we sought to determine the upstream kinase responsible for modulating RhoGEF2-EB1 binding. The iProteinDB database, which contains the phosphoproteomic data from *Drosophila* embryos, predicted AMPK would phosphorylate the serines proximal to the RhoGEF2 SKIP motif^43^. AMPK is best known for its ubiquitous role in nutrient sensing, but has recently been shown to regulate polarity and migration in a variety of cell types^44^. In *Drosophila*, AMPK regulates epithelial polarity in the embryo and can phosphorylate myosin II, providing an alternate pathway to activate contractility^45^. To determine if AMPK can phosphorylate RhoGEF2, we performed an in vitro phosphorylation assay with purified AMPK holoenzyme and an N terminal GST-RhoGEF2 fragment containing the SKIP motif and the nearby serines of interest (AA 1-164). A phosphorylation specific staining revealed that AMPK readily phosphorylated GST- RhoGEF2 (1-164) specifically at the seven serines near the SKIP motif, as a GST- RhoGEF2 7A (1-164) fragment (seven serines changed to alanine) did not show a detectable change in signal intensity upon incubation with AMPK (Fig. 6l-m). Having confirmed AMPK can phosphorylate RhoGEF2 *in vitro,* we next asked whether removing AMPK would perturb PGC migration. We generated AMPK null germline clone embryos and rescued known somatic defects by driving a GFP-AMPK with *nullo*-Gal4. We observed a significant increase in the number of PGCs which failed to arrive at the gonads in stage 14 embryos (Fig. 6n-o), suggesting AMPK was necessary for PGC migration. Based on these results, we propose the following model- AMPK spatiotemporally controls RhoGEF2 activity to impart the tunable modulation of migration speed necessary for accurate homing *in vivo* (Fig. 7).

**Fig. 7.**
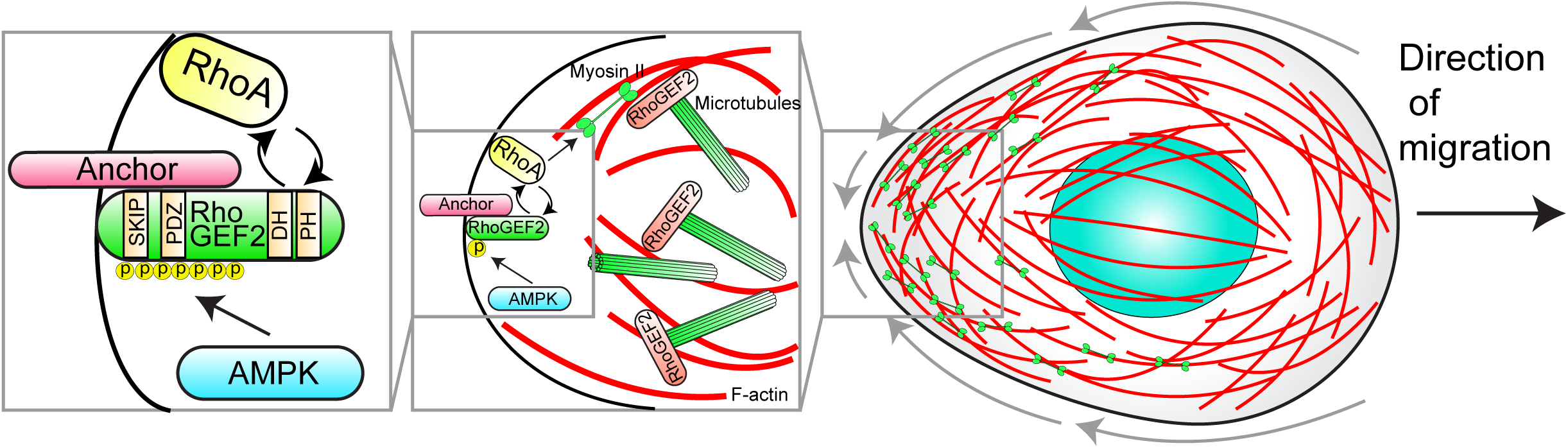
Model for RhoGEF2 cortical flow regulation in PGCs. Model for how RhoGEF2 modulates cortical flow and migration in PGCs. AMPK controls the free pool of RhoGEF2 by phosphorylating serines proximal to the SKIP motif to inhibit EB1 binding. Free RhoGEF2 accumulates on the membrane by binding an unknown anchor with its PDZ domain and triggers RhoA activation with its basal activity. Nascent RhoA activation allows for the recruitment of more RhoGEF2 through a postive feedback via binding the RhoGEF2 PH domain, generating robust RhoA activation. RhoGEF2 activity levels alter the actomyosin contractility gradient in a cell to organize and drive cortical flow. This cortical flow module allows PGCs to control their migration speed and orientation for accurate guidance in vivo.

## Discussion

Many cells possess the latent potential to adopt cortical flow driven amoeboid migration within confined, non-adhesive environments that promote contractility^7^. The hypothesized *in vivo* function of this migration strategy is to allow directional migration away from a contractile stimulus^4^, such as a wound, and/or to rapidly escape confinement through nuclear mechanosensation^9, 15^. We show that *Drosophila* PGCs endogenously utilize cortical actin flow to navigate *in vivo*, suggesting this migration mode is more widespread than previously appreciated. Moreover, while PGCs experience a confined cellular environment *in vivo*, we strikingly find that PGCs also maintain cortical flows *in vitro* without confinement and serum stimulation (Fig. 1e-f), suggesting cell intrinsic properties, such as basal actomyosin contractility, are themselves sufficient. Other factors which may contribute to the global flows we observe include the physical properties of the PGC actin cortex itself, as work in the *C. elegans* zygote has indicated that cortical viscosity can determine whether a local contraction induces a long range flow^12^. Cortical architecture may also be a determinant, as cortical connectivity influences the length scale of flow propagation^46, 47^. Indeed, elegant *in vitro* studies using reconstituted actin cortices have established that a percolation threshold exists, such that intermediate levels of crosslinking are required for flow propagation across the entire network^48^. Future studies need to determine the cortical properties and architecture that enables global cortical flow in PGCs.

PGC cluster dispersal and transepithelial migration depend on the GPCR Tre1^20, 21, 23–25^, which radially orients migration outwards, implying that GPCR signaling can orient cortical flow. How this mechanistically occurs is unclear, but it is likely to be distinct from the detailed pathways outlined in neutrophils and Dictyostelium, which follow the protrusive front and contractile rear paradigm. Indeed, we have previously shown that active CDC- 42 and Rac are not polarized in PGC clusters^20^ and that overexpression of constitutively active or dominant negative CDC42 does not impair migration^21^. Others have also noted that PI(3,4,5)P3, a phosphoinositide important for orienting protrusion driven cells, is similarly not polarized^25^. One common strategy to orient flow is to locally inhibit contractility, thereby creating an actomyosin contractility gradient that drives flow toward the opposite end of the cell. In both the *C. elegans* zygote and *Xenopus* oocyte, local inhibition occurs proximal to the microtubule-organizing center, suggesting a role for microtubules in this process, although this remains unclear in *C. elegans*^49, 50^. If PGCs follow a similar strategy, GPCR signaling may specify an axis of motility by determining which regions should not be the “rear”.

Our work further establishes a RhoGEF regulatory network responsible for tuning cortical flow (Fig. 7), whereby the PDZ dependent local accumulation of RhoGEF2 triggers nascent sites of RhoA activity which are further amplified by a PH domain dependent positive feedback loop. When RhoGEF2 is depleted, cortical actin flows are slower and more disorganized (Fig. 3j-n), suggesting RhoGEF2 establishes an actomyosin contractility gradient to both orient and induce sufficiently fast cortical flows for accurate migration. The RhoGEF network we uncover here does not require Gα_12/13_ and thus expands our knowledge of how RH-RhoGEFs are regulated in space and time. We speculate that this feedback loop could similarly be important for PDZ-RhoGEF, which operates at the rear of migrating neutrophils^51^, as Gα_12/13_ does not appreciably enhance its exchange activity^39^. As to why PGCs need to modulate their cortical flow speeds, PGCs need to stop once they reach their target and this could be accomplished by decreasing flow. Alternatively, PGCs may need to accelerate cortical flow to maintain their speed on different cellular substrates, such as when transitioning from an E-cadherin expressing endoderm to an N-cadherin expressing mesoderm. Such adaptive dynamics have been demonstrated in dendritic cells^52^.

Lastly, our findings expand the burgeoning role of AMPK in cell migration. We found that AMPK directly phosphorylates RhoGEF2 near its N-terminal SKIP motif, liberating it from EB1 dependent inhibition. Such regulation has precedence, as AMPK has previously been shown to phosphorylate CLIP-170 to reduce its affinity for microtubules^44^. Disrupting CLIP-170 phosphoregulation perturbs migration^44^, as we have similarly observed for RhoGEF2 in PGCs. In the context of directed PGC migration, local AMPK activation could orient cortical flow by releasing RhoGEF2 to establish a zone of high actomyosin contractility.

To conclude, the role of cortical flow driven amoeboid *in vivo* has been proposed to be a specialized response to external stimuli to enable rapid, context dependent persistent motility^4^. We find that this is the default mode by which Drosophila PGCs move and that it is receptive to external guidance, thematically similar to what has been observed in fat body cells^53^. Thus this could be a widespread single cell migration strategy *in vivo* because it can arise from the internal state of a cell, irrespective of its environment.

## Materials and methods

### Fly strains

All fly strains were maintained at 25°C, with experimental genotypes listed in Table S2. w^1118^ was used as the negative control. Transgenic lines were produced by Bestgene Inc. utilizing phiC31 integrase-mediated transgenesis. The landing sites utilized in this study were su(Hw)attP8 on the X chromosome, attP40 and su(HW)attP5 on the second chromosome, and attP2, VK27, VK28, and VK33 on the third chromosome.

### Constructs

Infusion (Clontech) cloning was utilized to create all constructs and Q5® High Fidelity DNA polymerase (NEB) was used for PCR. The *nos* and UAS with *nos*TCE-*pgc* 3’UTR (allows PGC targeting) backbones have previously been described^20^. pJFRC19 (Gerald Rubin^54^, Addgene 26224) was the backbone for LexAop driven constructs. pJRFC19 was modified to permit efficient, targeted expression in PGCs by removing the hsp70 promoter and SV40 3’UTR with a restriction digest and replacing them with the p-element promoter from pwalium22 and *nos*TCE-*pgc* 3’UTR, respectively. The *squash* (myosin II RLC in *Drosophila*) promoter driven RhoGEF2 constructs utilized pWALIUM22 (Perrimon lab^55^) as a backbone. The UAS sites and K10 3’UTR were replaced via PCR with the *squash* promoter and *squash* 3’UTR from pBS-Squ-mCherry (Eric Wieschaus^56^, Addgene 20163). The RhoGEF2 ORF was obtained from the DGRC (SD04476). All mutations were introduced with a site directed mutagenesis (NEB, Q5® site directed mutagenesis kit- E0554S). pGEX6P1-N-HA (Andrew Jackson and Martin Reijns, Addgene 119756) was utilized as the backbone for recombinant protein expression *E. coli*. All constructs were sequence verified before sending for injection.

#### LexAop2-p-degradFP- *nos*TCE-*pgc* 3’UTR

The degradFP system (Markus Affolter^26^, Addgene- 35579) was amplified by PCR and inserted into PJFRC19 containing the p-element promoter and *nos*TCE-*pgc* 3’UTR as described above. Fly lines were generated on su(Hw)attP8, attP40, and attP2.

#### Mat-tub LexA-GAD

Three fragments were cloned into pWALIUM22 with UAS sites and K10 3’UTR removed- (1) *αTub67C* promoter and 5’UTR were amplified via PCR from genomic DNA from P{matα4-GAL-VP16}67; P{matα4-GAL-VP16}15 females (BL 80361) , (2) nlsLexA::GADfl ORF was amplified from pBPnlsLexA::GADflUw (Gerald Rubin^54^, Addgene-26232), and (3) *αTub84B* 3’UTR was ordered as a Gblock from IDT technologies. Fly lines were generated on attP40 and attP2.

#### UASp-Gα_12/13_-Q303L-*nos*TCE-*pgc* 3’UTR

The Gα_12/13_(known as concertina in *Drosophila*) ORF (DGRC-LD04530) was amplified via PCR and cloned into pWALIUM22 containing the *nos*TCE-*pgc* 3’UTR described previously. Site directed mutagenesis was then utilized to generate the Q303L mutation. Fly lines were generated on attp40 and attP2.

#### nos-myosin-II-tdTomato-P2A-tdKatushka2-CAAX

*Nos* regulator elements drive the myosin II RLC (Eric Wieschaus^56^, Addgene 20163) fused to tdTomato (Michael Davidson, Addgene 54653), a P2A peptide, and tdKatushka2 (Michael Davidson^57^, Addgene 56041) with a CAAX box from human KRAS for membrane targeting. Three fragments were amplified via PCR and cloned into pWALIUM22 with *nos* regulatory elements described previously. The P2A and CAAX sequences were added to tdKatushka2 via primer. Fly lines were generated on attP40 and su(HW)attP5 on the second chromosome and attP2 and VK27 on the third chromosome.

#### squ-mNeongreen-RhoGEF2 constructs

For all constructs, mNeongreen (allele biotech) and RhoGEF2 (DGRC-SD04476) were first subcloned into pUC19 and sequence verified before subsequent cloning as a single fragment into pWALIUM22 with *squash* regulatory elements described above. For squ-mNeongreen-RhoGEF2-ΔPH RBD, the Anillin RhoA-GTP binding domain (RBD) was subcloned from the nos-tdTomato-Anillin-RBD-P2A-tdKatushka2-CAAX transgene generated previously^58^. Fly lines were generated on attP2, VK27, VK28, or VK33 on the third chromosome.

#### pAc EB1-mScarlet

EB1 (DGRC-RE41364) and mScarlet (Dorus Gadella^59^, Addgene 85042) were amplified with PCR and cloned into Ac5-Stable2-neo (Rosa Barrio and James Sutherland^60^, Addgene 32426) with inserts removed by PCR.

#### pAc mNeongreen RhoGEF2 (1-619) constructs

These constructs were subcloned via PCR from the corresponding squ-mNeongreen-RhoGEF2 full length constructs into Ac5-Stable2-neo (Rose Barrio and James Sutherland^60^, Addgene 32426).

#### pGEX-RG2-164, pGEX-RG2-164 7A

These constructs were subcloned via PCR from the corresponding squ-mNeongreen-RhoGEF2 full length constructs into pGEX6P1-N-HA (Andrew Jackson and Martin Reijns, Addgene 119756).

#### pGEX-Rho1

The Rho1 ORF (DGRC LD03419) was cloned into pGEX6P1-N-HA (Andrew Jackson and Martin Reijns, Addgene 119756).

### Live imaging

Embryos were produced at 25°C. For live imaging experiments from embryos, embryos were first dechorionated in 50% bleach for 3 minutes, extensively washed, collected onto a nylon mesh, and placed onto apple juice agar plates for visual staging. Appropriately staged embryos were subsequently oriented with their dorsal surface facing up with a fine needle, adhered to #1.5 glass coverslips (Thermo Fisher Scientific, 12-544-BP) with heptane glue, overlaid with halocarbon oil 27 (Sigma, H8733), and placed onto a gas permeable membrane (YSI, 098094). Live embryo imaging was performed on a custom Prairie Ultima (Bruker technologies) utilizing a Nikon CFI Apo IR 60x 1.27 NA water objective, a Chameleon Discovery tunable femtosecond laser with total power control (TPC), four external GaAsp PMTs (Hamamatsu 7422PA-40) , and driven by Prairie View 5.0 software.

A four channel upper non-descanned detector module allowed for simultaneous acquisition of four channels. An initial dichroic (t560lpxr) split the emission to two custom filter cubes (1) bandpass filters ET575/50m-2p and ET660/60M-2p with a T612LPXR-UF1 dichroic to simultaneously detect the emission from the fluorescent proteins tdTomato and tdKatushka2 used in this study and (2) bandpass filters ET460/50m-2p and ET525/50m-2p with a T495lpxr dichroic to detect the emission from the fluorescent proteins sfGFP and mNeongreen used in this study. The fourth channel, for detecting emission from fluorescent proteins such as CFP, was not utilized in this study. A 1080 nm wavelength was used for experiments where PGCs expressed reporters with tdTomato and tdKatushka2, while a 950 nm wavelength was utilized when PGCs co-expressed a given sfGFP or mNeongreen tagged transgene. All three fluorescent proteins (sfGFP/mNeongreen, tdTomato, and tdKatushka2) could be detected under sufficiently strong laser power at 950 nm. All filters were purchased from Chroma Technology Corp.

Live imaging of extracted PGCs and insect S2 cells *in vitro* was performed on a Nikon W1 spinning disk confocal microscope with a SR HP Plan Apo 100X 1.35 NA objective and Andor 888 EMCCD cameras driven by Nikon Elements software. The experimental setup is described below.

### PGC purification and experimentation

We capitalized on the early formation of PGCs before somatic cellularization is complete to develop an efficient protocol to specifically extract PGCs. We based our protocol off an existing protocol for FACS sorting *Drosophila* embryonic cells^61^. Embryos were first aged en masse to stage 5 of embryogenesis, homogenized in a dounce homogenizer in Schneiders medium (Thermofisher, 21720024), filtered through a 100 μm mesh to separate large embryonic fragments, centrifuged (500g for 1 minute), exchanged into hemolymph-like buffer (25 mM KCl, 90 mM NaCl, 4.8 mM NaHCO3, 80 mM d-glucose, 5 mM trehalose, 10 mM Hepes, PH 6.9) with .25% trypsin, and incubated in a 37°C water bath for 10 minutes to break up any cell clumps. The trypsin was then neutralized by adding Schneider’s medium with 10% FBS, the cell suspension was centrifuged, and the medium was exchanged for Schneiders’s medium with 1% (w/v) BSA.

For experiments without compression, PGCs were seeded directly onto a 8 well #1.5 glass lab-tek slide (Nunc) and allowed to settle for 1 hour before imaging. For under agarose experiments, a polydimethylsiloxane (PDMS) stencil was first created by curing Sylgard 184 (Dow) at a 1:10 crosslinker to polymer ratio and punching a central hole with a 14 mm punch. The stencil was then aligned and placed over the well of a 14 mm #1.5 glass bottom dish (MatTek). A 1% (w/v) ultrapure agarose (Thermofisher, 16500100) in PBS solution was then placed within the stencil and allowed to cure at room temperature. After the gel solidified, one end of the gel was gently picked up with a forcep, the PGC cell suspension was pipetted into the well, and the gel was gently placed back down. Schneiders medium was than pipetted around the gel. For experiments with Cyotochalasin D (Sigma, C8273) and Y-27632 (Millipore Sigma, 688001), the drugs were diluted in Schneiders medium, added around the gel in the dish, and allowed to incubate at room temperature for at least 1 hour before imaging.

### Particle image velicometry (PIV) analysis

All PIV analysis was performed in Matlab R2021a (Mathworks) using PIVLab^62–64^ with the following analysis settings-(1) Image pre-processing-CLAHE window size 10 pixels, (2) PIV settings-FFT window deformation, Pass 1- 64 pixels, step-32 pixels, Pass 2- 32 pixels, step-16 pixels, Pass 3- 16 pixels, step-8 pixels. Sub-pixel estimator- Gauss 2×3- point. Correlation robustness- Standard, (3) Post-processing- Velocity based validation- Standard deviation filter- 8, Local median filter- 3, Image based validation- Filter low contrast- this parameter was tuned between .005 to .02 depending on a given cell.

All results were exported to Matlab and subsequently analyzed and visualized using custom written code. For average cortical flow speed, the average vector magnitude for each xy vector across all time points was averaged. For cumulative distributions, all vector magnitudes at each xy position across all time points aggregated. For cosine similarity analysis, the cosine similarity between vectors at each xy position at consecutive time points was computed with the following formula-

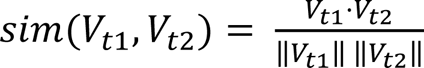

The computed cosine similarities were than averaged to obtain the mean value for each cell.

### Image processing and analysis

Two photon images presented in figures and movies were denoised using the CANDLE^65^ package for Matlab in Matlab 2016a (Mathworks). The settings utilized were beta = .3, patch radius = 2, and search radius = 2. All quantitative analysis was performed on raw data. For insect S2 or PGC *in vitro* imaging, a 1 pixel Gaussian filter was applied for denoising.

To quantify actin cortical flow *in vivo*, kymographs were produced for 5 actin clusters per cell and used to estimate flow speed. The mean of the 5 clusters was taken as the actin flow speed. Cell speed was computed by creating a kymograph with the membrane images. To quantify myosin II cortical flow *in vivo,* a kymograph was used to estimate the flow speed of a single myosin II foci. Cell speed was calculated as described above.

To quantify the period of circular flows in PGCs, a kymograph was generated across the cell from the myosin II image and the resulting distance between three high intensity regions, indicating flow passing through this region, was used to calculate the period. We defined cells as exhibiting periodic cortical flows if the flows traveled around the circumference of the cell at least three times during imaging.

For cell tracking and quantification of cell speed in ImageJ and Matlab, regions of interest (ROI) were first defined in ImageJ for cell segmentation. ROIs were subsequently imported into Matlab and cells were automatically segmented from ROIs using the “func_threshold” function^66^ from all Z planes. The cell centroid was calculated as the average XYZ pixel value for all segmented pixels. Only cells which exhibited a persistence (minimal distance between initial and endpoint/total distance traveled) > .5 were used in analysis. All other analysis was completed using a combination of ImageJ and custom written scripts in Matlab described below.

Quantification of polarized myosin II in cells has been previously described^20^. Briefly, regions of high myosin II (at least >=1.2 fold greater intensity than cytoplasm) along with a corresponding region in the cytoplasm were manually segmented in ImageJ and imported into Matlab. The computed ratios between these ROIs was taken as the polarized intensity. Quantification of polarized myosin II orientation with respect to the endoderm center in individual cells has been previously described^20^. Briefly, the angle between the following line segments was used to calculate orientation- (1) PGC nuclei and region of polarized myosin II (defined above) and (2) PGC nuclei and endoderm center.

To quantify PGC dispersal rate, PGCs trans-migrating across the endoderm were manually annotated from timelapse image stacks.

Quantification of fluorescence intensity as a function of distance from cluster centroid has previously been described^20^. Briefly, individual PGC clusters were manually segmented in ImageJ with the PGC specific membrane marker (tdKatushka2-CAAX) along with a second ROI in the background for normalization. The ROIs were then imported into Matlab and used as masks for the other fluorescent channels of interest. Pixel intensities were then placed into 50 equally spaced bins according to distance from the cluster centroid.

Quantification of posterior myosin II intensity over time has previously been described^20^. Briefly, individual PGCs were manually segmented in ImageJ in the Z plane which contained the greatest polarized myosin II intensity. ROIs were subsequently imported into Matlab and segmented PGCs were computationally rotated vertically with posterior towards the bottom and posterior was defined as the lowest 20% of segmented rows. The mean intensity in this posterior region over all time points was used as posterior myosin II intensity.

To quantify sfGFP-RhoGEF2 degradation, an ROI was defined in the central plane of the PGC cluster using the PGC membrane marker (tdKatushka2-CAAX) in ImageJ. This ROI was then used as a mask for the sfGFP-RhoGEF2 channel and the mean intensity of the segmented pixels was quantified.

To quantify phospho-myosin II intensity in fixed S2 cells as a function of mNeongreen RhoGEF2 construct expression, maximum intensity projections were first generated from Z stacks taken from S2 cells expressing a given mNeongreen RhoGEF2 construct and stained for phospho-myosin II. Each cell was manually segmented using the mNeongreen signal with an ROI to obtain the mean construct expression level and this ROI was subsequently used as a mask to generate the mean phospo-myosin II intensity. Each individual cell’s construct expression level and corresponding mean phospho-myosin II intensity were then placed into three equally spaced bins (low, medium, and high). The bins were equivalent across experimental conditions in the same plots.

To quantify cellularization furrow intensity in stage 5 embryos, ten somatic furrow tip intensities were calculated by manually segmentation from PGC membrane marker images (this reporter has low expression in somatic cells) and normalized to the intensity of the top of the furrow, near the apical surface of the cell.

To quantify immunoblot or gel images, ROIs were defined in ImageJ around each band and the integrated density was extracted and normalized to the loading control or input.

### Cell culture and transfection

S2 cells were cultured in Schneider’s medium with 10% (v/v) fetal bovine serum (ThermoFisher, 16140071) and 1% (v/v) penicillin streptomycin (ThermoFisher, 15140122). Effectene (Qiagen, 301425) was utilized for all transfections following manufacturer’s recommendations. Transfected S2 cells were plated onto Labtek slides (ThermoFisher, 155409) coated with 50 μg/ml Concanavalin A (Cayman Chemical, 14951) diluted in PBS for 1 hour at room temperature for live imaging. For immunofluorescence, S2 cells were seeded onto 16 mm circular Concanavalin A coated coverslips (same as above) for at least 1 hour before proceeding to immunostaining.

### Antibodies

Primary antibodies utilized in this study for immunofluorescence- rabbit anti-vasa (1:5000, R. Lehmann), guinea pig anti-phospho myosin II (1:1000, R. Ward), chicken anti-vasa (1:500, R. Lehmann), and rabbit anti-RhoGEF2 (1:2500, J. Grosshans).

Secondary antibodies utilized in this study for immunofluorescence- Cy3 AffiniPure Donkey Anti-Rabbit IgG (Jackson Immunoresearch, 711-165-152), Cy3 AffiniPure Donkey Anti-Guinea Pig IgG (Jackson Immunoresearch, 706-165-148), Alexa Fluor 488 AffiniPure Donkey Anti-Rabbit IgG (Jackson Immunoresarch, 711-545-152), and Alexa Fluor® 488 AffiniPure Donkey Anti-Chicken IgY (Jackson Immunoresearch, 703-545-155).

Primary antibodies utilized for immunoblotting- mouse anti-mNeongreen (1:1000, Chromotek-32F6), rabbit anti-mNeongreen (1:1000, Cell Signaling-53061), mouse anti-α-tubulin (1:4000, Sigma-T6199), mouse anti-RFP (1:1000, Chromotek-6G6), and rabbit anti-GST (1:2000, Cell Signaling-2622).

Secondary antibodies utilized in this study for immunoblotting-HRP Goat anti-rabbit IgG (1:10000, Abcam-ab6721) and HRP Rabbit anti-mouse IgG (1:10000, Abcam-ab6728).

### Immunofluorescence

Embryos were first dechorionated in 50% bleach for 3 minutes, extensively washed, collected on a nylon mesh, and transferred to a scintillation vial containing a 1:1 (v/v) mixture of heptane and 4% paraformaldehyde (Electron Microscopy Sciences, 15714-S) in PBS on a shaker for 20 minutes. The paraformaldehyde was subsequently removed with a Pasteur pipette and replaced with methanol and the scintillation vial was vigorously shaked by hand for 30 seconds to remove the vitelline membrane. Embryos were kept in methanol at -20°C until subsequent processing. Embryos stored in methanol were gradually rehydrated with PBST (.3% Triton X-100 (Sigma, T8787)) and blocked in PBST with 1% bovine serum albumin (BSA) (Sigma, A4503) for 60 minutes at room temperature. All primary antibodies were diluted in PBST with 1% BSA and applied overnight at 4°C. After extensive washing, appropriate secondary antibodies (1:500, Jackson Immunoresearch) were diluted in PBST with 1% BSA and incubated with samples for 3 hours at room temperature. Embryos were washed and subsequently mounted in Vectashield (Vector Laboratories, H-1000) and imaged with a Zeiss LSM 800 using Zen Blue 2.3 with a 20X .8 NA air objective using a pinhole size of 1 AU.

Insect S2 cells were fixed in 4% paraformaldehyde, briefly permeabilized with .1% Triton X-100, washed, and blocked overnight in 5% (w/v) BSA in PBS. Primary and secondary antibodies, diluted in 5% BSA in PBS, were than applied at room temperature for 1 hour with extensive washes between these steps. The samples were then mounted with Prolong Diamond (Molecular Probes, P36965) onto glass slides and imaged with a Nikon W1 spinning disk confocal microscope with an Apo 60x 1.40 NA oil objective and Andor 888 EMCCD cameras driven by Nikon Elements software.

### In vitro translation

A TNT T7 Quick Coupled Transcription/Translation System (Promega, L1170) with Transcend tRNA (Promega, L5061) was utilized to *in vitro* translate the RhoGEF2 PH domain according to the manufacturer’s recommendation.

### Co-immunoprecipitation

S2 cell lysates were harvested 3-4 days after transfection in co-IP buffer (50 mM Tris-HCl pH 7.5, 250 mM NaCl, 1 mM EDTA, 1 mM MgCl2, .2% NP-40, 10% glycerol) with protease/phosphatase inhibitors (Thermo Fisher Scientific, 78442), incubated on ice for 30 minutes, clarified by centrifugation, and measured with a BCA assay (Thermo Fisher Scientific, 23225). Equal amounts of lysate (1 mg) were then immunoprecipitated with mNeongreen-trap magnetic agarose resin (Chromotek, ntma) for 1 hour at 4°C, washed, and then eluted with LDS buffer (Thermo Fisher Scientific, NP0007).

### GST protein purification

pGEX-Rho1, pGEX-RG2-164, pGEX-RG2-164 7A, and pGEX6P1-N-HA were transformed into BL21(DE3) competent cells (New England Biolabs, C2527H) and grown overnight at 37°C in a starter culture. The starter cultures were then diluted 1:200 into 1 L cultures, shaken for ∼1.5 hours at 37°C until reaching an OD600 of .6-.8, induced with 100 μM IPTG, and shaken for 20-24 hours at room temperature. The bacterial cultures were then pelleted by centrifugation, washed once with PBS, and flash frozen and stored at -80°C until further processing. To harvest recombinant GST proteins, frozen bacteria pellets were lysed in lysis buffer (20 mM HEPES (pH. 7.5), 150 mM NaCl, 5 mM MgCl2, 1 % triton, 1 mM DTT with protease/phosphatase inhibitors), sonicated in a Bioruptor Pico (Diagenode), and clarified by centrifugation at 4°C. The clarified lysate was then incubated with Glutathione Sepharose 4B resin (Millipore Sigma, GE17-0756-01) for 2 hours at 4°C, washed, and then resuspended in storage buffer (HBS, 5mM MgCl2, 1 mM DTT) with 33% glycerol.

### In vitro kinase assay

GST-RG2-164 and GST-RG2-164 7A were eluted from Glutathione Sepharose 4B resin with elution buffer (50 mM Tris-CL PH 8, 30 mM glutathione) and concentrated and exchanged into HEPES-Brij buffer (50 mM Na-HEPES (pH 7.4), 1 mM DTT, 5 mM MgCl2, and 0.02% Brij-35) with an Amicon Ultra-2 10K NMWL filtration unit (UFC201024). Protein concentrations were then estimated by SDS-PAGE with BSA standards followed by InstantBlue Coomassie stain (ab119211). In vitro kinase assays were performed with ∼2 μg of substrate, 50 ng of AMPK holoenzyme (EMD Millipore, 14-840), .2 mM ATP (EMD Millipore, 1191-5GM), and .3 mM AMP (EMD Millipore, 118110-5GM) where indicated for 30 min at 30°C. Reactions were terminated by adding LDS buffer and samples were separated by SDS-PAGE. Phosphorylated proteins were detected with a ProQ diamond stain (Thermo Fisher Scientific, P33302) using a modified protocol^67^ and total proteins were subsequently detected with InstantBlue Coomassie stain.

### In vitro binding assay with GST-Rho1

10 μg GST-Rho1 per reaction was first resuspended in nucleotide exchange buffer (50 mM HEPES (PH 7.08), 5 mM EDTA, .1 mM EGTA, 50 mM NaCL, .1 mM DTT) and loaded with 0.5 mM GDP (Sigma-Aldrich, G7127) or GTPγS (Cytoskeleton, bs01) for 30 min at 30°C. The reaction was terminated by adding 20 mM MgCl2. GST-Rho1-GDP and GST-Rho1-GTPγS were then exchanged into HEPES-LS buffer (20 mM HEPES, pH 7.5, 150 mM NaCl, 10% glycerol, 0.1% Triton-X-100) and incubated with ∼1 μg (estimated by SDS-PAGE) of *in vitro* translated RG2-PH domain for 1 hour at 4°C. Following washes, the remaining bound products were eluted with LDS buffer.

### Protein extraction from embryos

Overnight embryo collections were dechorionated in 50% bleach for 3 minutes and transferred with a paint brush to ice cold RIPA buffer (Thermo Scientific, 89900) with protease/phosphatase inhibitors in a 1.5 ml Eppendorf tube. The embryos were subsequently homogenized with a motorized pestle and incubated with periodic mixing for 30 minutes on ice. The lysates were then clarified by centrifugation and protein concentrations were measured with a BCA assay for subsequent SDS-PAGE.

### SDS-PAGE and immunoblotting

Equal amounts of protein were separated via SDS PAGE using a 1.5 cm 4-12% Bis-Tris gel (Thermo Fisher Scientific, NP0336) or 1 cm 4-12% Bis-Tris gel for ProQ diamond stain with 1x MOPS running buffer (Thermo Fisher Scientific, NP0001), transferred onto a PVDF membrane (EMD Millipore, IPFL07810), blocked for 1 hour in 2% (v/v) BSA (Bioworld, 40220068-1) in TBST (TBS with .3% Tween-20) and probed overnight with primary antibodies diluted in the above blocking buffer. Following extensive washes, HRP linked ECL conjugated secondary antibodies, Goat anti-rabbit IgG and Rabbit anti-mouse IgG, were used to identify proteins of interest and were visualized with a Supersignal West Pico PLUS ECL substrate (Thermo Fisher Scientific, 34579).

For *in vitro* binding assays with GST-Rho1, in vitro translated RhoGEF2 PH domains were detected on immunoblots with streptavidin-HRP (1:5000, Thermo Fisher Scientific, SA10001) and visualized with a Supersignal West Pico PLUS ECL substrate (Thermo Fisher Scientific, 34579). The membranes were subsequently incubated with hydrogen peroxide to quench the bound HRP, washed, and subject to GST immunoblotting using the above protocol.

### Statistics

All experiments were performed with at least 2 independent replicates and the number of embryos/cells analyzed are noted in the figures. All statistical comparisons were carried out in Prism (GraphPad) using Mann Whitney tests for pair-wise comparisons.

## Data Availability

All data is available from the corresponding authors upon reasonable request.

## Author Contributions

B.L. designed experiments, performed research, analyzed data, and wrote the manuscript with help from J.L. on experimentation and data analysis. R.L. supervised the project and wrote the manuscript.

## Acknowledgements

We acknowledge the FlyBase team for maintaining the Drosophila database (funded by NHGRI [ U41HG000739 ]) and the Drosophila Genomics Resource Center, supported by NIH grant 2P40OD010949, for reagents. Stocks obtained from the Bloomington Drosophila Stock Center (NIH P40OD018537) were used in this study. We thank Daniela Drummond-Barbosa for sharing fly lines, Robert Ward for sharing the phospho-myosin II antibody, and Jörg Großhans for sharing the RhoGEF2 antibody. We also thank Michael Cammer and the NYULH DART Microscopy Laboratory (P30CA016087) for microscopy support. This work is supported by NIH grant R37HD41900 to R.L and formerly by HHMI, where R.L. was an investigator. B.L. was a New York Stem Cell Foundation Druckenmiller Fellow.

## Competing interests

The authors declare no competing interests.

---

**Fig. S1.**
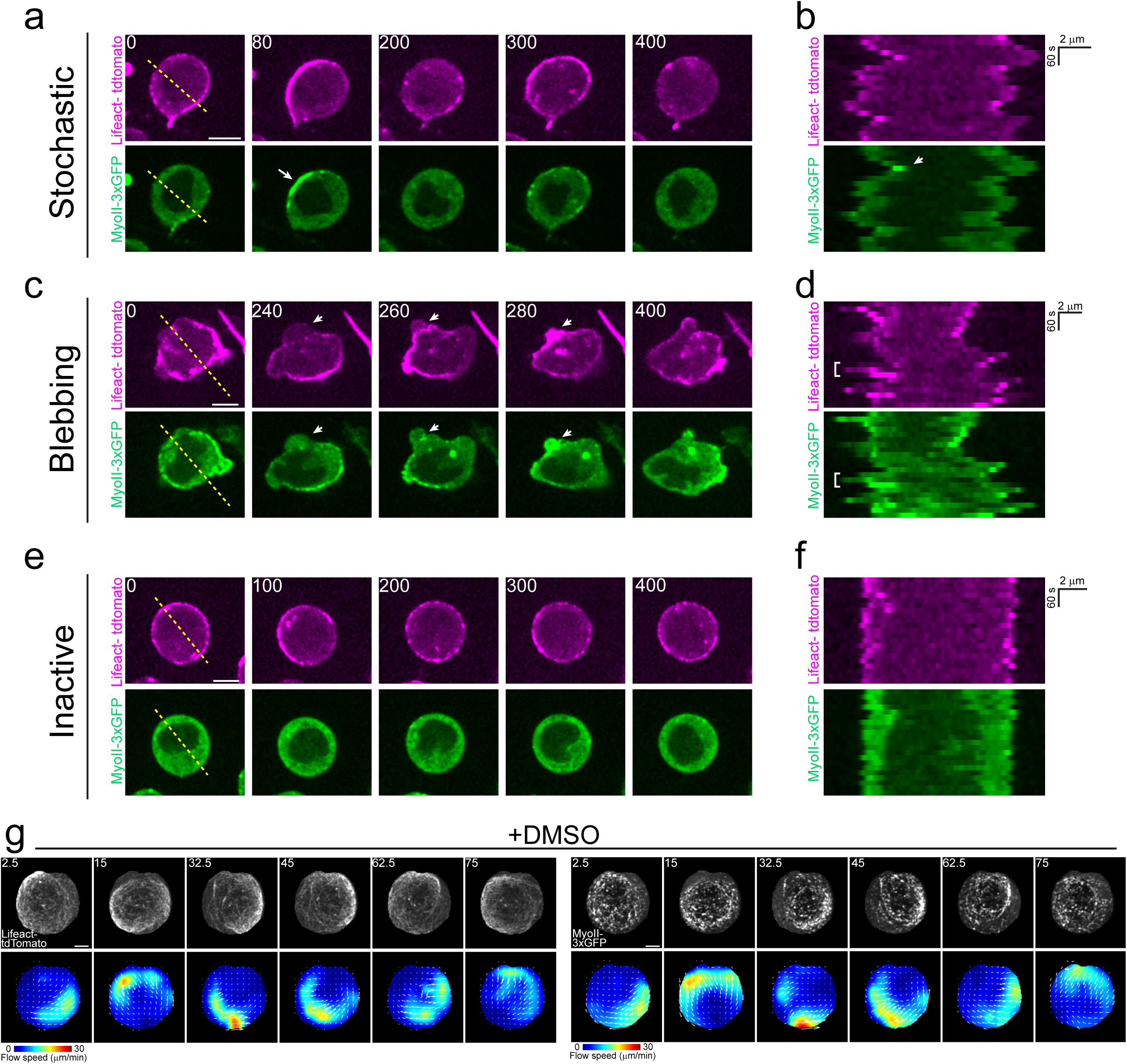
PGC dynamics *in vitro*. (**a-f**) Timelapse imaging of representative extracted PGCs expressing lifeact-tdTomato and myosin II-3xGFP on cover- glass which are stochastic (**a-b**), blebbing (**c-d**), or inactive (**e-f**). Yellow dotted lines indicate where the kymographs in **b**,**d**,**f** were taken. White arrow in **a** indicates a region where myosin II has accumulated. The equivalent region in the kymograph in **b** is also indicated with a white arrow. White arrow in **c** indicates a bleb and its subsequent retraction. Bleb emergence and retraction is highlighted by a white bracket in **d**. Scale bars, 10 μm. (**g**) Top- timelapse imaging of repre- sentative extracted PGC under agarose expressing lifeact-tdtomato (left panels) and myosin II-3xGFP (right panels) treated with DMSO. Bottom- PIV flow analysis between the image above and the previous time point. Flow speed is color coded with the indicated color bar. White arrows are the flow vectors at the indicated positions scaled to flow magnitude. Scale bar, 5 μm. All times are in seconds.

**Fig. S2.**
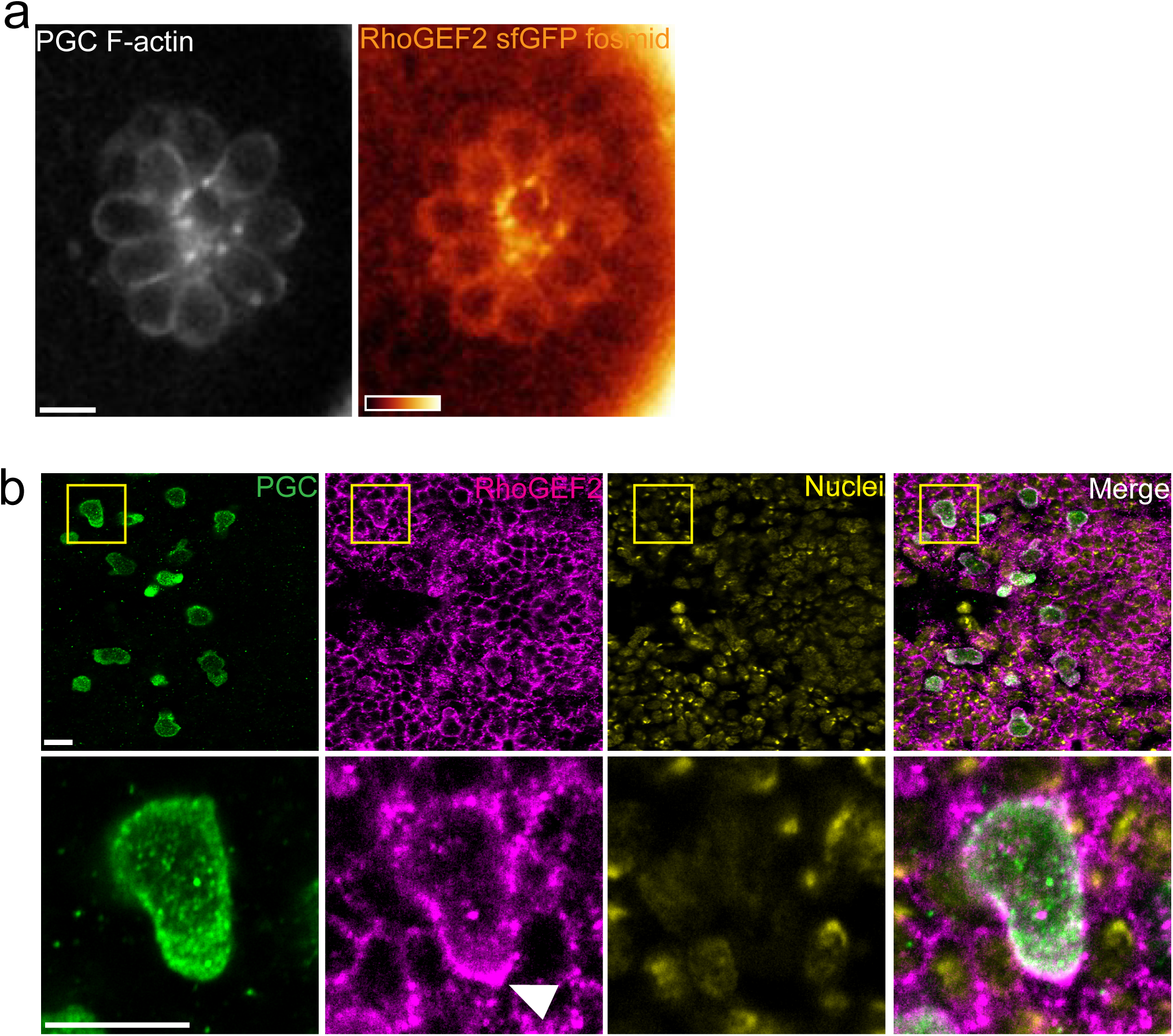
RhoGEF2 localization under endogenous regulation in PGCs. (**a)** Representative two photon image from a central plane in a PGC cluster expressing lifeact-tdTomato, tdKatush- ka2-CAAX (not shown), and a RhoGEF2-sfGFP fosmid. The image is representative of 9 embryos imaged. (**b**) Representative immunofluorescent image from a stage 11 embryo, where PGCs are migrating towards the mesoderm (top and bottom of this image). The yellow box in top images show the enlarged image below. The white arrow indicates where RhoGEF2 has accumulated at the rear of the PGC. This image is representative of 5 embryos imaged. Scale bars, 10 μm in all images.

**Fig. S3.**
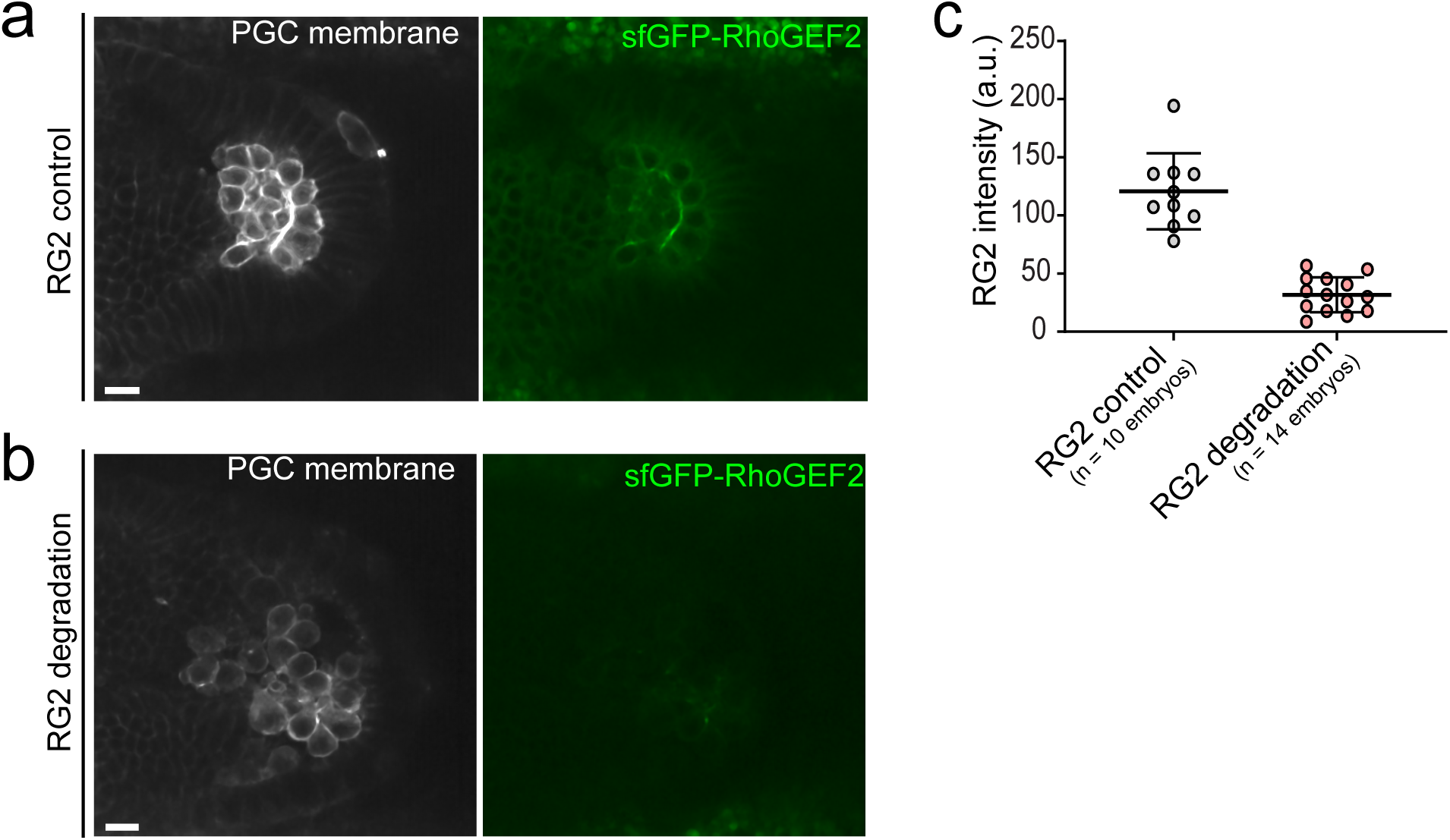
Quantification of sfGFP-RhoGEF2 degradation with degradFP. (**a-b)** Representative two photon image from a central plane in a PGC cluster expressing tdkatushka2-CAAX (PGC membrane) and sfGFP-RhoGEF2 under control (**a**) and RhoGEF2 degradation (**b**) conditions. The image is representa- tive of n=10 embryos (RG2 control) and n=14 embryos (RG2 degradation). RhoGEF2, RG2. Scale bars, 10 μm. (**b**) Quantification of mean RhoGEF2 intensity in the central plane of PGC clusters under the indicated conditions. The number of embryos imaged is indicated.

**Fig. S4.**
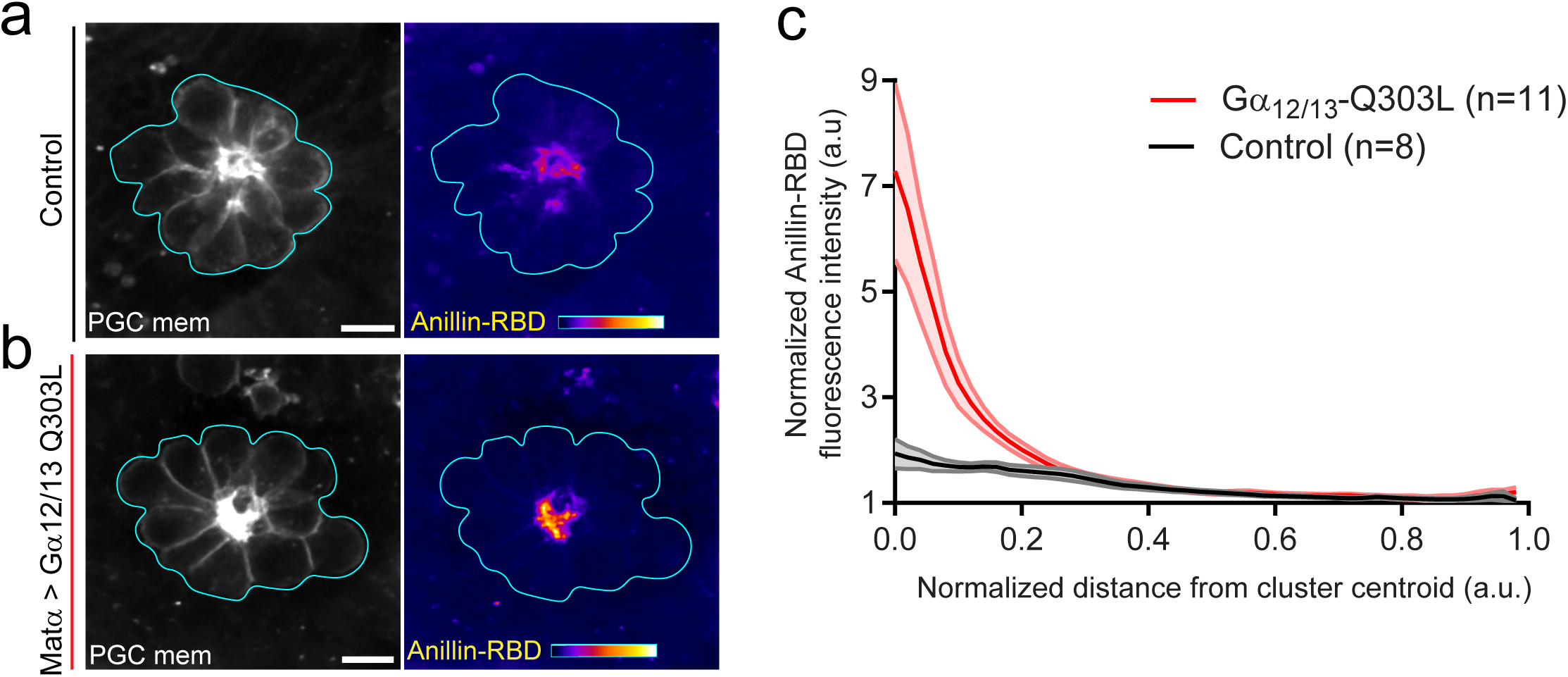
Expression of constitutively active Gα_12/13_ enhances active RhoA polarity. (**a-b)** Representative two photon image from a central plane in a PGC cluster expressing Anillin-RBD-tdTomato and tdkatushka2-CAAX under the indicated conditions. RhoA binding domain, RBD. The image is representative of n=8 embryos (control) and n=11 embryos (Gα12/13-Q303L). Scale bars, 10 μm. (**b**) Quantification Anillin-RBD-tdTomato intensity as a function of distance from the cluster centroid. The number of embryos analyzed is indicated. Error bars are SEM.

**Fig. S5.**
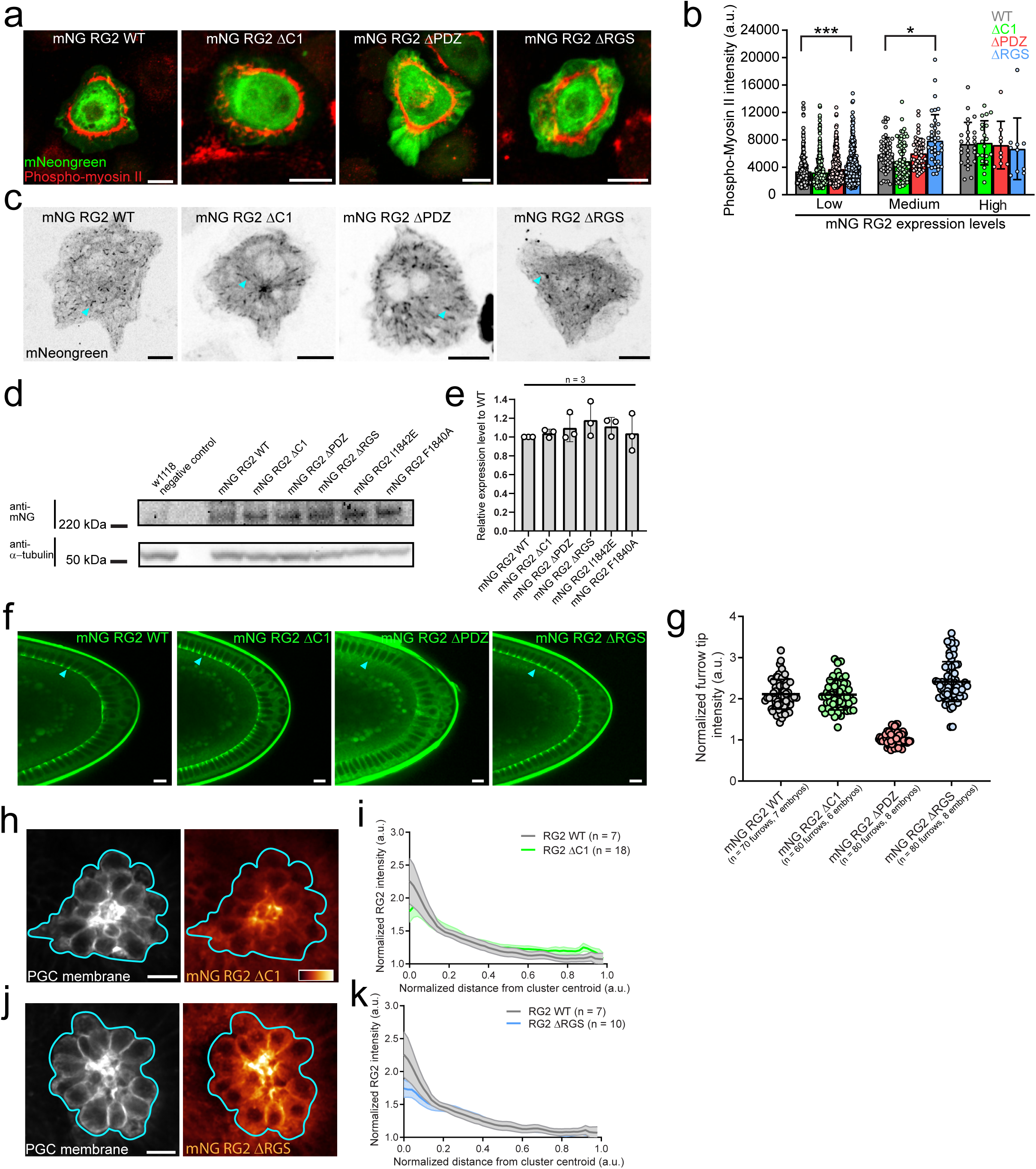
Characterization of RhoGEF2 domain truncation constructs. **(a**) Representative immunofluorescence images from insect S2 cells overexpressing the indicated mNeongreen RhoGEF2 construct and co-stained with a phospho-myosin II antibody. Note that microtubule plus-end tracking is abolished under the utilized fixation conditions. Scale bars, 10 μm. (**b**) Quantification of phospho-myosin II intensity as a function of mNeongreen RhoGEF2 construct expression level. Expression levels were split into 3 equally sized bins. The number of cells analyzed from 3 independent experi- ments is- WT low- 317, WT medium- 50, and WT high- 26, ΔC1 low- 517, ΔC1 medium- 74, and ΔC1 high- 20. ΔPDZ low- 410, ΔPDZ medium- 47, and ΔPDZ high- 9. ΔRGS low- 259, ΔRGS medium- 38, and ΔRGS high- 10. (**c**) Representative image from live S2 cells expressing the indicated mNeongreen RhoGEF2 construct. The cyan arrows indicate that microtubule plus-end tracking is intact in all constructs. (**d)** Representative immunoblot from embryonic lysates expressing the indicated mNeongreen RhoGEF2 transgene stained with the indicated antibodies. w1118 is a negative control for antibody specificity. (**e**) Quantification of mNeongreen RhoGEF2 transgene expression levels relative to WT. The number of experiments is indicated. ((**f**) Representative two photon image from the posterior pole of stage 5 embryos expressing the indicated mNeongreen RhoGEF2 transgene. Cyan arrows inicate mNeongreen RhoGEF2 transgene enrichment on furrow tips, with the exception of the ΔPDZ transgene. (**g**) Quantification of furrow tip enrichment of different mNG RhoGEF2 transgenes. The number of furrows and embryos analyzed are indicated. Error bars are S.D. (**h,j**) Repre- sentative two photon image from central plane of PGC clusters expressing tdKatushka2 CAAX (PGC membrane marker) and the indicated mNeongreen RhoGEF2 transgene. mNeongreen RhoGEF2 is pseudocolored with the color bar in the image. (**i,k**) Quantifi- cation of mNeongreen RhoGEF2 transgene intensity as a function of distance from the cluster centroid. The number of embryos analyzed is indicated. Error bars are SEM. Scale bars, 10 μm in all images. *=p<.05, *** = p<.001.

**Fig. S6.**
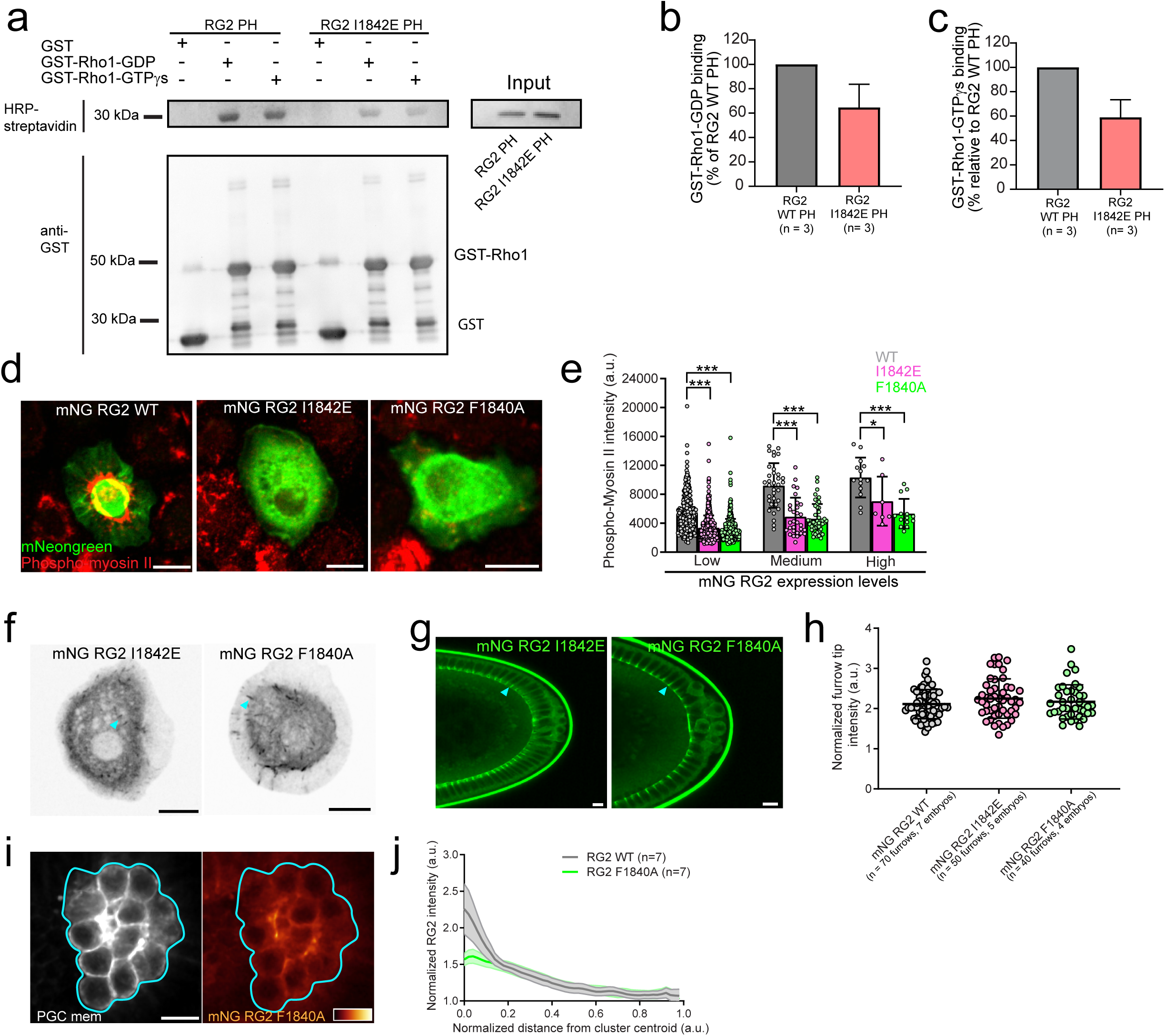
Characterization of RhoGEF2 PH mutants. (**a)** Representative immunoblot from an in vitro binding assay between the in vitro translated, biotin conjugated, RhoGE2 PH domains and purified GST fusion proteins. The top image is stained with HRP-streptavidin to identify the amount of biotin-conjugated RhoGEF2 PH domain pulled down with each GST protein. The same membrane was then stripped and stained with anti-GST antibody to assess if equal amounts of GST protein were present in each sample (bottom). (**b-c**) Quantification of the amount of RhoGEF2 PH domain bound to GST-Rho1-GDP (**b**) or GST-Rho1-GTPγs (**c**) relative to the WT PH domain. (**d**) Representative immunofluorescence images from insect S2 cells overexpressing the indicated mNeongreen RhoGEF2 construct and co-stained with a phospho-myosin II antibody. Note that microtubule plus-end tracking is abolished under the utilized fixation conditions. (**e**) Quantifi- cation of phospho-myosin II intensity as a function of mNeongreen RhoGEF2 construct expression level. Expression levels were split into 3 equally sized bins. The number of cells analyzed from 3 independent experiments is- WT low- 403, WT medium- 34, and WT high- 14, 1842E low- 573, 1842E medium- 30, and 1842E high- 7. F1840A low- 501, F1840A medium- 47, and F1840A high- 14. (**f**) Representative image from live S2 cells expressing the indicated mNeongreen RhoGEF2 construct. Cyan arrow indicates that microtubule plus-end tracking is intact in all constructs. (**g**) Representative two photon image from the posterior pole of stage 5 embryos expressing the indicated mNeongreen RhoGEF2 transgene. Cyan arrows inicate mNeongreen RhoGEF2 transgene enrich- ment on furrow tips. (**h**) Quantification of furrow tip enrichment of different RhoGEF2 transgenes. The number of furrows and embry- os analyzed are indicated. Error bars are S.D. (**i**) Representative two photon image from central plane of PGC cluster expressing tdKatushka2 CAAX (PGC membrane marker) and the mNeongreen RhoGEF2 F1840A transgene. mNeongreen RhoGEF2 is pseudocolor with the color bar in the image. (**j**) Quantification of mNeongreen RhoGEF2 F1840A intensity as a function of distance from the cluster centroid compared to WT. The number of embryos analyzed is indicated. Error bars are SEM. Scale bars, 10 μm in all images. *=p<.05, *** = p<.001.

**Fig. S7.**
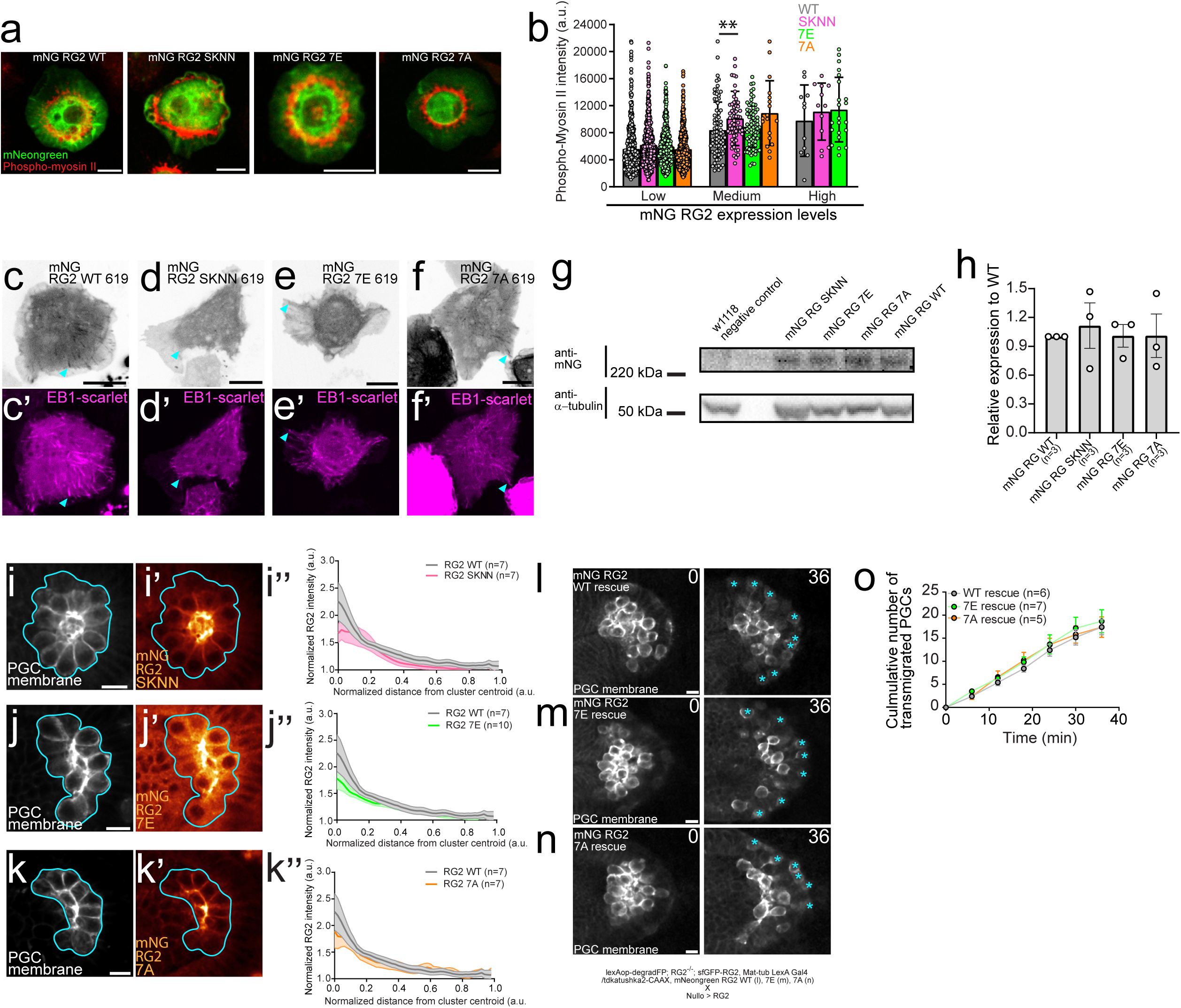
Characterization of RhoGEF2 SKIP mutants. (**a)** Representative immunofluorescence images from insect S2 cells overexpressing the indicated mNeongreen RhoGEF2 construct and co-stained with a phospho-myosin II antibody. Note that microtubule plus-end tracking is abolished under the utilized fixation conditions. (**b**) Quantificatoin of phospho-myosin II intensity as a function of mNeongreen RhoGEF2 construct expression level. Expression levels were split into 3 equally sized bins. The number of cells analyzed from 3 independent experiments is- WT low- 474, WT medium- 76, and WT high- 11, SKNN low- 573, SKNN medium- 59, and SKNN high- 12. 7E low- 450, 7E medium- 74, and 7E high- 22. 7A low- 606 and 7A medium- 18. We did not observe any high expressing mNeongreen RhoGEF2 7A S2 cells. (**c-f**) Repre- sentative images from live S2 cells co-expressing a given truncated mNeongreen RhoGEF2 construct (1-619) and EB1-mScarlet. Cyan arrow indicates that microtubule plus-end tracking is maintained in **c,f** but abolished in **d,e**. (**g)** Representative immunoblot from embryonic lysates expressing the indicated mNeongreen RhoGEF2 transgenes stained with the indicated antibodies. w1118 is a negative control for antibody specificity. (**h**) Quantification of mNeongreen RhoGEF2 transgene expression levels relative to WT. The number of experiments is indicated. (**i-k**) Representative two photon image from central plane of a PGC cluster expressing tdKatush- ka2 CAAX (PGC membrane marker) and the indicated mNeongreen RhoGEF2 transgene. mNeongreen RhoGEF2 is pseudocolored with the color bar in the image. (**i’’,k’’, k’’**) Quantification of mNeongreen RhoGEF2 transgene intensity as a function of distance from the cluster centroid. The number of embryos analyzed is indicated. Error bars are SEM. (**l-n**) Representative two photon timelapse imaging of PGC clusters expressing tdKatushka2-CAAX (PGC membrane marker) dispersing under the indicated rescue conditions. Genotypes are indicated below. Cyan asterisks mark transmigrated PGCs. Times are in minutes. (**o**) Quantification of the total number of transmigrated PGCs over time under the indicated resuce conditions. The number of embryos analyzed is noted. Error bars are SEM. Scale bars, 10 μm in all images. ** = p<.01.

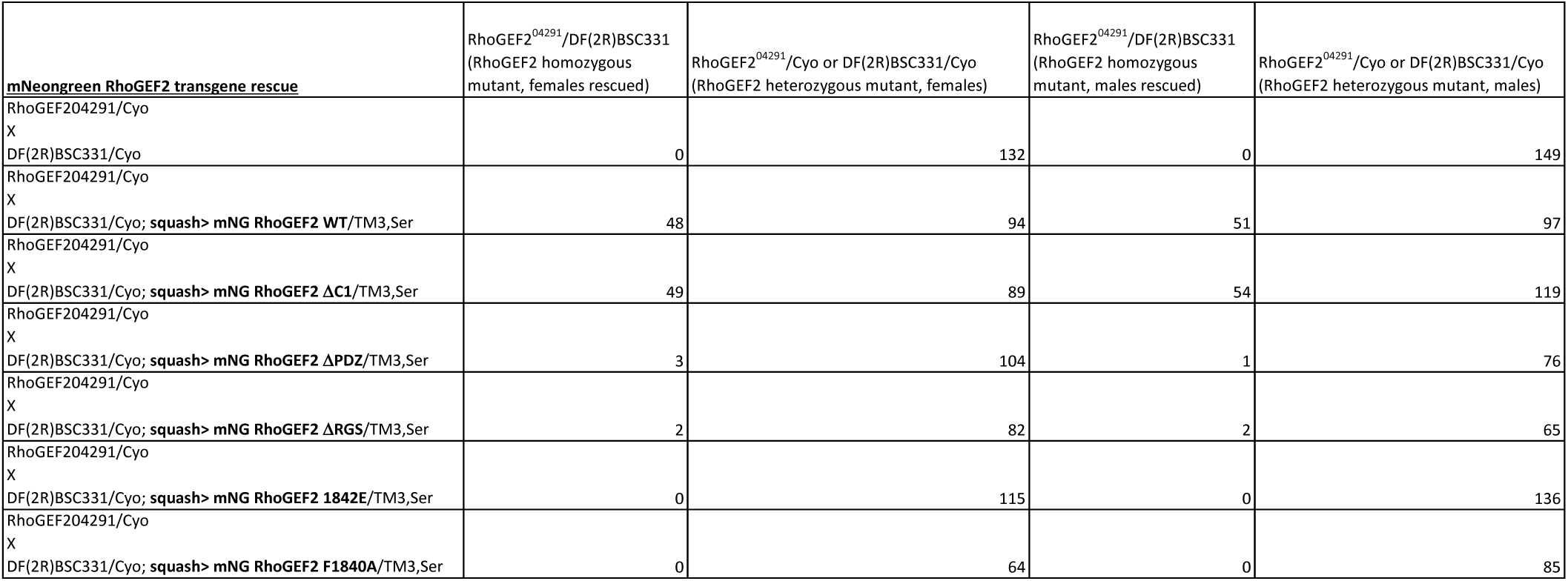

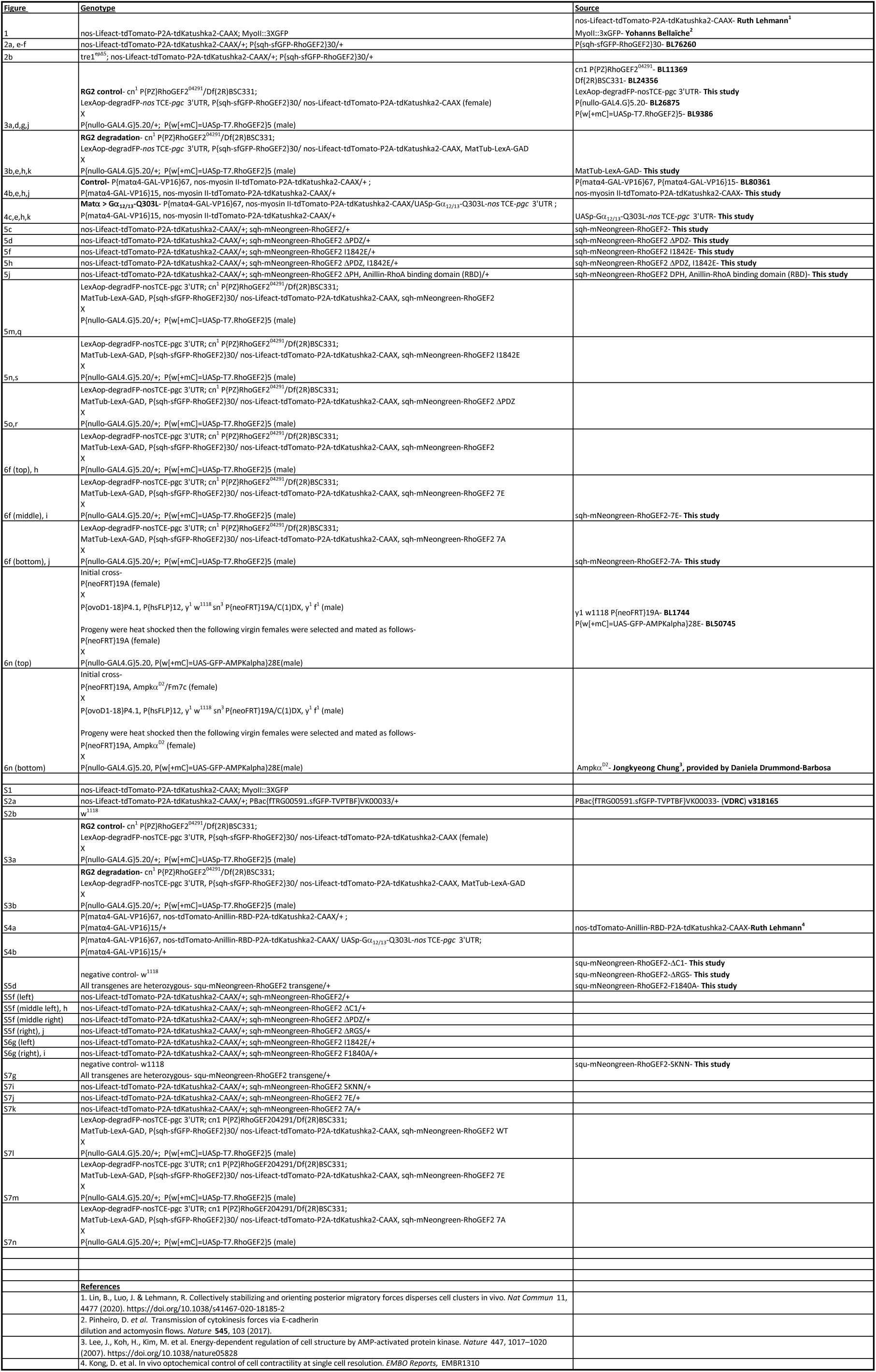

## Notes

### Competing Interest Statement

The authors have declared no competing interest.

## References

1. Case, L. B. & Waterman, C. M. Integration of actin dynamics and cell adhesion by a three- dimensional, mechanosensitive molecular clutch. Nature Cell Biology 17, 955–963, doi:10.1038/ncb3191 (2015).

2. Bergert, M. et al. Force transmission during adhesion-independent migration. Nature Cell Biology 17, 524–529, doi:10.1038/ncb3134 (2015).

3. Poincloux, R. et al. Contractility of the cell rear drives invasion of breast tumor cells in 3D Matrigel. 108, 1943–1948, doi:10.1073/pnas.1010396108 %J Proceedings of the National Academy of Sciences (2011).

4. Ruprecht, V. et al. Cortical Contractility Triggers a Stochastic Switch to Fast Amoeboid Cell Motility. Cell 160, 673–685, doi:10.1016/j.cell.2015.01.008 (2015).

5. Venturini, V. et al. The nucleus measures shape changes for cellular proprioception to control dynamic cell behavior. Science 370, eaba2644, doi:10.1126/science.aba2644 (2020).

6. Logue, J. S. et al. Erk regulation of actin capping and bundling by Eps8 promotes cortex tension and leader bleb-based migration. eLife 4, e08314, doi:10.7554/eLife.08314 (2015).

7. Liu, Y.-J. et al. Confinement and Low Adhesion Induce Fast Amoeboid Migration of Slow Mesenchymal Cells. Cell 160, 659–672, doi:10.1016/j.cell.2015.01.007 (2015).

8. Shih, W. & Yamada, S. Myosin IIA Dependent Retrograde Flow Drives 3D Cell Migration. Biophysical Journal 98, L29–L31, doi:https://doi.org/10.1016/j.bpj.2010.02.028 (2010).

9. Lomakin, A. J. et al. The nucleus acts as a ruler tailoring cell responses to spatial constraints. Science 370, eaba2894, doi:10.1126/science.aba2894 (2020).

10. Kardash, E. et al. A role for Rho GTPases and cell–cell adhesion in single-cell motility in vivo. Nature Cell Biology 12, 47–53, doi:10.1038/ncb2003 (2010).

11. Tozluoğlu, M. et al. Matrix geometry determines optimal cancer cell migration strategy and modulates response to interventions. Nature Cell Biology 15, 751–762, doi:10.1038/ncb2775 (2013).

12. Mayer, M., Depken, M., Bois, J. S., Jülicher, F. & Grill, S. W. Anisotropies in cortical tension reveal the physical basis of polarizing cortical flows. Nature 467, 617–621, doi:10.1038/nature09376 (2010).

13. Maiuri, P. et al. Actin Flows Mediate a Universal Coupling between Cell Speed and Cell Persistence. Cell 161, 374–386, doi:https://doi.org/10.1016/j.cell.2015.01.056 (2015).

14. Callan-Jones, A. C. & Voituriez, R. Actin flows in cell migration: from locomotion and polarity to trajectories. Current Opinion in Cell Biology 38, 12–17, doi:https://doi.org/10.1016/j.ceb.2016.01.003 (2016).

15. Ullo, M. F. & Logue, J. S. ADF and cofilin-1 collaborate to promote cortical actin flow and the leader bleb-based migration of confined cells. eLife 10, e67856, doi:10.7554/eLife.67856 (2021).

16. Barton, L. J., LeBlanc, M. G. & Lehmann, R. Finding their way: themes in germ cell migration. Current Opinion in Cell Biology 42, 128–137, doi:https://doi.org/10.1016/j.ceb.2016.07.007 (2016).

17. Richardson, B. E. & Lehmann, R. Mechanisms guiding primordial germ cell migration: strategies from different organisms. Nature Reviews Molecular Cell Biology 11, 37–49, doi:10.1038/nrm2815 (2010).

18. Blaser, H. et al. Migration of Zebrafish Primordial Germ Cells: A Role for Myosin Contraction and Cytoplasmic Flow. Developmental Cell 11, 613–627, doi:https://doi.org/10.1016/j.devcel.2006.09.023 (2006).

19. Olguin-Olguin, A. et al. Chemokine-biased robust self-organizing polarization of migrating cells in vivo. 118, e2018480118, doi:10.1073/pnas.2018480118 %J Proceedings of the National Academy of Sciences (2021).

20. Lin, B., Luo, J. & Lehmann, R. Collectively stabilizing and orienting posterior migratory forces disperses cell clusters in vivo. Nature Communications 11, 4477, doi:10.1038/s41467-020-18185-2 (2020).

21. Kunwar, P. S., Starz-Gaiano, M., Bainton, R. J., Heberlein, U. & Lehmann, R. Tre1, a G protein- coupled receptor, directs transepithelial migration of Drosophila germ cells. PLoS Biol 1, E80–E80, doi:10.1371/journal.pbio.0000080 (2003).

22. Hons, M. et al. Chemokines and integrins independently tune actin flow and substrate friction during intranodal migration of T cells. Nature Immunology 19, 606–616, doi:10.1038/s41590-018-0109-z (2018).

23. Kunwar , P. S. et al. Tre1 GPCR initiates germ cell transepithelial migration by regulating Drosophila melanogaster E-cadherin. Journal of Cell Biology 183, 157–168, doi:10.1083/jcb.200807049 %J Journal of Cell Biology (2008).

24. LeBlanc, M. G. & Lehmann, R. Domain-specific control of germ cell polarity and migration by multifunction Tre1 GPCR. Journal of Cell Biology 216, 2945–2958, doi:10.1083/jcb.201612053 %J Journal of Cell Biology (2017).

25. Kim, J. H., Hanlon, C. D., Vohra, S., Devreotes, P. N. & Andrew, D. J. Hedgehog signaling and Tre1 regulate actin dynamics through PI(4,5)P2 to direct migration of <em>Drosophila</em> embryonic germ cells. Cell Reports 34, doi:10.1016/j.celrep.2021.108799 (2021).

26. Caussinus, E., Kanca, O. & Affolter, M. Fluorescent fusion protein knockout mediated by anti- GFP nanobody. Nature Structural & Molecular Biology 19, 117–121, doi:10.1038/nsmb.2180 (2012).

27. Großhans, J. r. et al. RhoGEF2 and the formin Dia control the formation of the furrow canal by directed actin assembly during Drosophila cellularisation. Development 132, 1009–1020, doi:10.1242/dev.01669 %J Development (2005).

28. Barrett, K., Leptin, M. & Settleman, J. The Rho GTPase and a Putative RhoGEF Mediate a Signaling Pathway for the Cell Shape Changes in Drosophila Gastrulation. Cell 91, 905–915, doi:10.1016/S0092-8674(00)80482-1 (1997).

29. Häcker, U. & Perrimon, N. DRhoGEF2 encodes a member of the Dbl family of oncogenes and controls cell shape changes during gastrulation in Drosophila. Genes Dev 12, 274–284, doi:10.1101/gad.12.2.274 (1998).

30. Dawes-Hoang, R. E. et al. folded gastrulation, cell shape change and the control of myosin localization. Development 132, 4165–4178, doi:10.1242/dev.01938 %J Development (2005).

31. Parks, S. & Wieschaus, E. The drosophila gastrulation gene concertina encodes a Gα-like protein. Cell 64, 447–458, doi:https://doi.org/10.1016/0092-8674(91)90652-F (1991).

32. Manning, A. J., Peters, K. A., Peifer, M. & Rogers, S. L. Regulation of epithelial morphogenesis by the G protein-coupled receptor mist and its ligand fog. Sci Signal 6, ra98–ra98, doi:10.1126/scisignal.2004427 (2013).

33. Fuse, N., Yu, F. & Hirose, S. Gprk2 adjusts Fog signaling to organize cell movements in Drosophila gastrulation. Development 140, 4246–4255, doi:10.1242/dev.093625 %J Development (2013).

34. Kölsch, V., Seher, T., Fernandez-Ballester, G. J., Serrano, L. & Leptin, M. Control of Drosophila Gastrulation by Apical Localization of Adherens Junctions and RhoGEF2. 315, 384–386, doi:doi:10.1126/science.1134833 (2007).

35. Wenzl, C., Yan, S., Laupsien, P. & Großhans, J. Localization of RhoGEF2 during Drosophila cellularization is developmentally controlled by slam. Mechanisms of Development 127, 371–384, doi:https://doi.org/10.1016/j.mod.2010.01.001 (2010).

36. Stein, J. A., Broihier, H. T., Moore, L. A. & Lehmann, R. Slow as Molasses is required for polarized membrane growth and germ cell migration in Drosophila. Development 129, 3925–3934, doi:10.1242/dev.129.16.3925 %J Development (2002).

37. Kunwar, P. S., Starz-Gaiano, M., Bainton, R. J., Heberlein, U. & Lehmann, R. Tre1, a G Protein- Coupled Receptor, Directs Transepithelial Migration of Drosophila Germ Cells. PLOS Biology 1, e80, doi:10.1371/journal.pbio.0000080 (2003).

38. Medina, F. et al. Activated RhoA is a positive feedback regulator of the Lbc family of Rho guanine nucleotide exchange factor proteins. J Biol Chem 288, 11325–11333, doi:10.1074/jbc.M113.450056 (2013).

39. Wells, C. D. et al. Mechanisms for Reversible Regulation between G13 and Rho Exchange Factors*. Journal of Biological Chemistry 277, 1174–1181, doi:https://doi.org/10.1074/jbc.M105274200 (2002).

40. Rich, A., Fehon, R. G. & Glotzer, M. Rho1 activation recapitulates early gastrulation events in the ventral, but not dorsal, epithelium of Drosophila embryos. eLife 9, e56893, doi:10.7554/eLife.56893 (2020).

41. Rogers, S. L., Wiedemann, U., Häcker, U., Turck, C. & Vale, R. D. Drosophila RhoGEF2 Associates with Microtubule Plus Ends in an EB1-Dependent Manner. Current Biology 14, 1827–1833, doi:https://doi.org/10.1016/j.cub.2004.09.078 (2004).

42. Honnappa, S. et al. An EB1-Binding Motif Acts as a Microtubule Tip Localization Signal. Cell 138, 366–376, doi:https://doi.org/10.1016/j.cell.2009.04.065 (2009).

43. Hu, Y. et al. iProteinDB: An Integrative Database of Drosophila Post-translational Modifications. G3 (Bethesda) 9, 1-11, doi:10.1534/g3.118.200637 (2019).

44. Nakano, A. et al. AMPK controls the speed of microtubule polymerization and directional cell migration through CLIP-170 phosphorylation. Nature Cell Biology 12, 583–590, doi:10.1038/ncb2060 (2010).

45. Lee, J. H. et al. Energy-dependent regulation of cell structure by AMP-activated protein kinase. Nature 447, 1017–1020, doi:10.1038/nature05828 (2007).

46. Koenderink, G. H. & Paluch, E. K. Architecture shapes contractility in actomyosin networks. Current Opinion in Cell Biology 50, 79–85, doi:https://doi.org/10.1016/j.ceb.2018.01.015 (2018).

47. Ding, W. Y. et al. Plastin increases cortical connectivity to facilitate robust polarization and timely cytokinesis. Journal of Cell Biology 216, 1371–1386, doi:10.1083/jcb.201603070 %J Journal of Cell Biology (2017).

48. Ennomani, H. et al. Architecture and Connectivity Govern Actin Network Contractility. Current Biology 26, 616–626, doi:https://doi.org/10.1016/j.cub.2015.12.069 (2016).

49. Benink, H. A., Mandato, C. A. & Bement, W. M. Analysis of cortical flow models in vivo. Mol Biol Cell 11, 2553–2563, doi:10.1091/mbc.11.8.2553 (2000).

50. Gan, W. J. & Motegi, F. Mechanochemical Control of Symmetry Breaking in the Caenorhabditis elegans Zygote. 8, doi:10.3389/fcell.2020.619869 (2021).

51. Wong, K., Van Keymeulen, A. & Bourne, H. R. PDZRhoGEF and myosin II localize RhoA activity to the back of polarizing neutrophil-like cells. J Cell Biol 179, 1141–1148, doi:10.1083/jcb.200706167 (2007).

52. Renkawitz, J. et al. Adaptive force transmission in amoeboid cell migration. Nature Cell Biology 11, 1438–1443, doi:10.1038/ncb1992 (2009).

53. Franz, A., Wood, W. & Martin, P. Fat Body Cells Are Motile and Actively Migrate to Wounds to Drive Repair and Prevent Infection. Developmental Cell 44, 460–470.e463, doi:https://doi.org/10.1016/j.devcel.2018.01.026 (2018).

54. Pfeiffer, B. D. et al. Refinement of Tools for Targeted Gene Expression in Drosophila. Genetics 186, 735–755, doi:10.1534/genetics.110.119917 (2010).

55. Ni, J.-Q. et al. Vector and parameters for targeted transgenic RNA interference in Drosophila melanogaster. Nature Methods 5, 49–51, doi:10.1038/nmeth1146 (2008).

56. Martin, A. C., Kaschube, M. & Wieschaus, E. F. Pulsed contractions of an actin–myosin network drive apical constriction. Nature 457, 495–499, doi:10.1038/nature07522 (2009).

57. Shcherbo, D. et al. Far-red fluorescent tags for protein imaging in living tissues. Biochemical Journal 418, 567–574, doi:10.1042/bj20081949 (2009).

58. Kong, D. et al. In vivo optochemical control of cell contractility at single-cell resolution. EMBO reports 20, e47755, doi:https://doi.org/10.15252/embr.201947755 (2019).

59. Bindels, D. S. et al. mScarlet: a bright monomeric red fluorescent protein for cellular imaging. Nature Methods 14, 53–56, doi:10.1038/nmeth.4074 (2017).

60. González, M. et al. Generation of stable Drosophila cell lines using multicistronic vectors. Sci Rep 1, 75–75, doi:10.1038/srep00075 (2011).

61. Defaye, A. & Perrin, L. in Hox Genes: Methods and Protocols (eds Yacine Graba & René Rezsohazy) 183–195 (Springer New York, 2014).

62. Thielicke, W. & Sonntag, R. Particle Image Velocimetry for MATLAB: Accuracy and enhanced algorithms in PIVlab. Journal of Open Research Software 9, doi:10.5334/jors.334 (2021).

63. Thielicke, W. & Stamhuis, E. J. PIVlab – Towards User-friendly, Affordable and Accurate Digital Particle Image Velocimetry in MATLAB. Journal of Open Research Software 2, doi:10.5334/jors.bl (2014).

64. Thielicke, W. The flapping flight of birds, [S.n.], (2014).

65. Coupé, P., Munz, M., Manjón, J. V., Ruthazer, E. S. & Louis Collins, D. A CANDLE for a deeper in vivo insight. Medical Image Analysis 16, 849–864, doi:https://doi.org/10.1016/j.media.2012.01.002 (2012).

66. Ridler, T. W. & Calvard, S. Picture Thresholding Using an Iterative Selection Method. *IEEE Transactions on Systems*, Man, and Cybernetics 8, 630–632, doi:10.1109/TSMC.1978.4310039 (1978).

67. Agrawal, G. K. & Thelen, J. J. in The Protein Protocols Handbook (ed John M. Walker) 579–585 (Humana Press, 2009).

